# CHK1-mediated regulation of TOP1 catalytic activity suppresses replication and transcription-associated genomic instability

**DOI:** 10.1101/2024.10.23.619942

**Authors:** Ananda Guha Majumdar, Nitish Chauhan, Pooja Gupta, Mahesh Subramanian, Birija Sankar Patro

**Author notes:** Corresponding Authors [ (MS); (BSP)].

## Abstract

The Topoisomerase 1 (TOP1) catalytic cycle involves a TOP1-DNA-covalent-complex (TOP1cc), which, if stabilized, can induce rapid accumulation of potentially lethal DNA double strand breaks (DSBs). Although TOP1cc are critically associated with genome instability, it is not yet precisely known how cells regulate TOP1cc level, under unperturbed physiological-condition, to prevent its accumulation and lethal consequences. We discovered a key role of CHK1 in phosphorylating TOP1 at Serine-320 and stimulating TOP1-catalytic cycle to minimise genome wide accumulation of TOP1cc in cancers. Pharmacological or genetic ablation of CHK1-mediated TOP1-phosphorylation leads to stalled replication forks and generates copious amounts of replication/transcription-associated DSBs, R-loops and transcription-replication collisions, eventually leading to chromosomal instability. Further, TOP1ccs stabilized due to CHK1 inhibition are not efficiently targeted by cellular TOP1cc-removal machineries. Since multiple patient clinical trials are ongoing with TOP1- and CHK1-targeting drugs, current finding of CHK1-mediated regulation of TOP1cc may help in better understanding the therapeutic outcomes.

## Introduction

Topoisomerase 1 (TOP1), an indispensable enzyme in humans, relaxes DNA supercoils during replication as well as transcription^1,2^. During its catalytic cycle, a transient TOP1-DNA covalent complex (TOP1cc) intermediate is formed, where the enzyme makes a phosphotyrosine bond involving the catalytic Tyrosine 723 and the 3’ end of the scissile site. Under ambient conditions, the TOP1-mediated religation step is significantly faster than the cleavage step, resulting in trace levels of TOP1ccs inside cells^1^. However, a number of conditions such as DNA damage of spontaneous origin or the presence of a TOP1 poison [*e.g*., camptothecins (CPT)] may interfere with the kinetics of the religation reaction, resulting in the stabilization of TOP1ccs. TOP1ccs, if stabilized, act as sources of double stranded breaks (DSBs) (due to conflicts with the replication or transcription machinery) as well RNA/DNA hybrids (R-loops)^1–3^. Hence, due to the potentially lethal outcomes of TOP1cc stabilization, the cellular machinery goes to great lengths to rapidly resolve such structures^3^. The cellular response to TOP1ccs may be broadly categorized into (a) those focussed on the removal of the covalently trapped protein and/or a DNA segment containing the same and (b) those revolving around prevention of conflicts with replication and/or the transcription machinery. Responses of the first category involve (i) proteasomal/non-proteasomal processing of TOP1ccs followed by their removal *via* TDP1/TDP2, (ii) nuclease mediated removal of a TOP1cc-containing oligonucleotide segment involving a battery of context-specific nucleases involving, but not limited to, the XPF-ERCC complex, Mus81/Eme1, MRE11, CtIP etc^4–6^. Recently, we have shown TDP1 and CtIP-mediated removal of TOP1cc is dependent on the Werner Syndrome (WRN) helicase^7^. Responses of the second category involve a myriad of processes, including PARP1, RECQL helicases and the Fanconi anemia pathway to name a few. These responses are mutually interdependent, and synchronised into a highly efficient and robust process which faithfully ensures genomic stability in the face of continuous genomic insults^8,9^. More recently, TOP1ccs have been reported to be eliminated through TEX264-driven selective autophagy, which also plays a critical role in repair of TOP1cc-associated DNA lesions^10^.

Post-translational modifications of TOP1 have been shown to play a pivotal role in regulation of TOP1 catalytic activity as well as removal of TOP1ccs. CPT-mediated TOP1 poisoning has been shown to trigger rapid DNA-PK mediated phosphorylation of TOP1 at Serine 10, which is essential for its proteasomal degradation^11^. Ubiquitination^12^, SUMOylation^13^ and NEDDylation^14^ have been shown to regulate proteasomal as well as non-proteasomal degradation of TOP1ccs^5,15^. PARylation, on the other hand, operates at multiple levels ranging from regulation of TOP1cc degradation to recruitment of TDP1 and fork remodelling factors, as well as regulation of nucleolar localization of TOP1^16–19^. Along with the role of post translational modifications in orchestrating cellular response to TOP1 poisoning, there has been significant interest in regulatory circuits of TOP1 catalytic activity under unperturbed conditions. The c-ABL kinase has been shown to regulate TOP1 catalytic activity through phosphorylation of Tyrosine 268 of TOP1^20^. CK-2 regulates TOP1 catalytic activity, CPT sensitivity and interaction with p14/ARF through phosphorylation of TOP1 Tyrosine 506^21,22^. Further, mitotic phosphorylation of TOP1 at Serine 10 (CK-2 mediated), Serine 21 (protein kinase Cα mediated), Serine 112 and Serine 394 (CDK1 mediated) has been shown to promote TOP1 catalytic activity^23^. CHK1, a master cell cycle checkpoint regulator and a prime DNA damage response-associated kinase plays a pivotal role in the downstream cellular events in the wake of TOP1 poisoning. Previously, we have shown that CHK1 is rapidly activated in response to TOP1ccs in a manner dependent on WRN helicase^7,24^. Post activation, CHK1 performs indispensable functions in the canonical DNA damage-induced enforcement of cell cycle checkpoints^25,26^. In spite of the relatively well understood accounts of the role played by CHK1 kinase in regulating cellular response to TOP1ccs, there have been no reports concerning any upstream role played by CHK1 in regulating the equilibrium of the TOP1 catalytic cycle under ambient conditions. In our current work, we have combined observations from cell biology, live cell imaging, *in silico* analysis as well as *in vitro* experiments to demonstrate the direct regulation of TOP1 catalytic activity inside cells by CHK1 kinase.

Our results indicate that pharmacological inhibition of CHK1 results in copious stabilization of TOP1ccs on the genome. CHK1 phosphorylates TOP1 *in vitro*; interestingly, Serine 320, An *in silico* predicted, but previously unreported CHK1 target site on TOP1 is indeed phosphorylated on catalytically engaged TOP1. Ablation of CHK1-mediated phosphorylation on TOP1 results in compromised catalytic activity of the enzyme, which is reflected in widespread TOP1cc stabilization and slower catalytic kinetics *in vitro*. Further, abolishing CHK1-mediated phosphorylation of TOP1 results in replication and transcription-associated DNA damage, chromosomal aberrations, transcription-replication collisions and accumulation of R-loops. Taken together, our results suggest a previously unreported role of basal cellular CHK1 kinase activity in suppression of genomic instability through regulation of the TOP1 catalytic cycle at physiological level.

## Results

### Screening of small molecule kinase inhibitors reveals CHK1 as a regulator of TOP1 dynamics

In order to discover novel regulators of TOP1 dynamics at physiological level, we performed a RADAR assay-based screen for TOP1cc stabilization (**Figure 1A**). We constructed a manually curated library of 25 small molecule kinase inhibitors (**Supplementary Table 1**). To ensure maximum coverage in terms of kinases while maintaining requisite specificity, we selected inhibitors which specifically target all (or majority) of the isoforms of a particular kinase with minimal experimentally reported cross-inhibition of other kinase families. The library was estimated to target a total of ∼42 cellular kinases which are known to regulate various aspects of the DNA damage response. The RADAR screen with U2-OS cells was carried out at 3 different concentrations of these kinase inhibitors at 3 different time points (2, 4, and 6 h) (**Figure 1A**). Our results revealed previously reported as well as novel hits. While previously reported regulators of TOP1 such as CDK1 and DNA-PK were successfully captured in the screen, CHK1, ATR and p38 were novel candidates (**Figure 1B-Z, S1A**).

**Figure 1:**
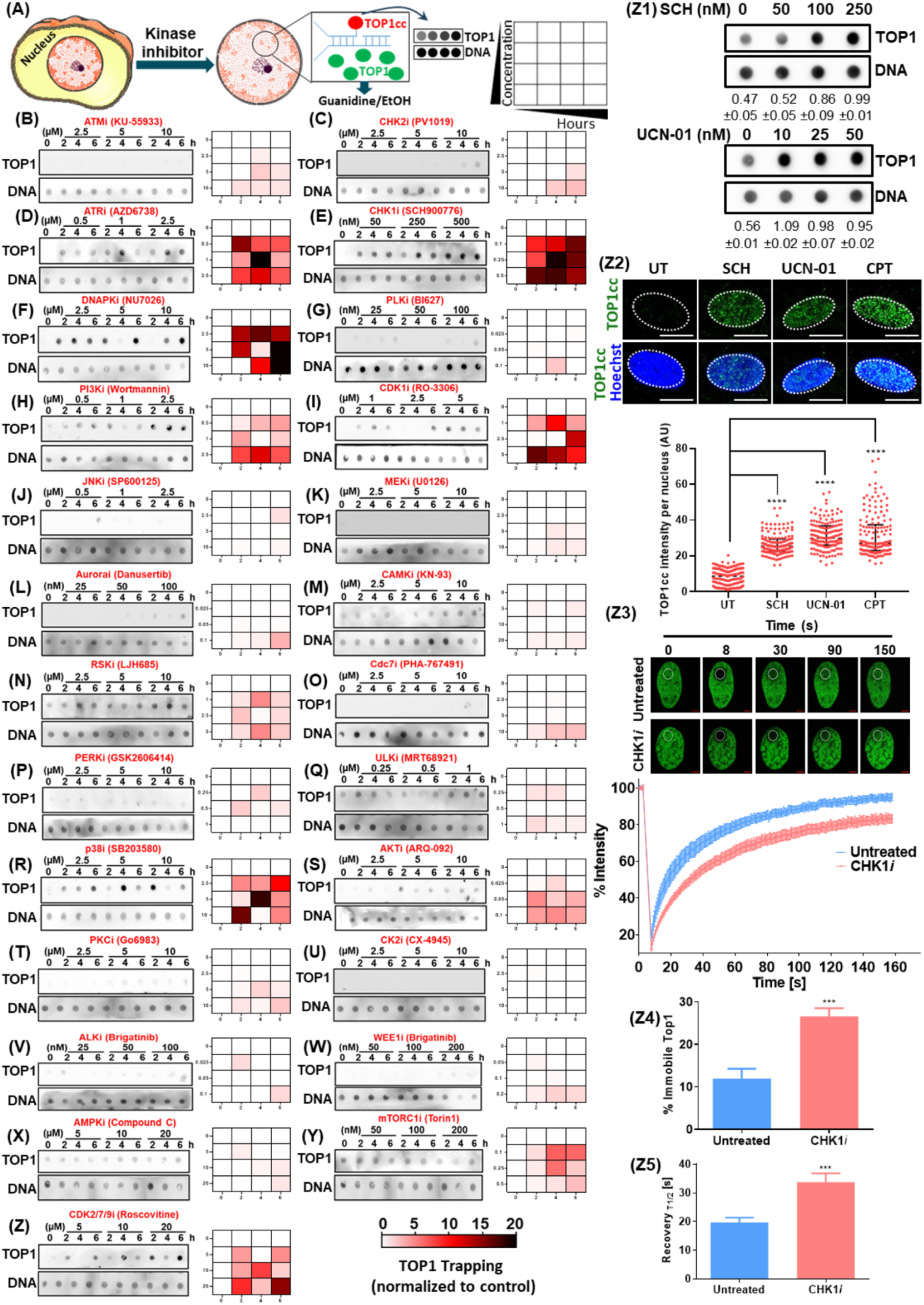
Pharmacological inhibition of CHK1 stabilizes TOP1ccs. (A) Experimental scheme of RADAR assay screen for TOP1cc stabilization by kinase inhibitors. (B-Z) Dot blots and corresponding heat maps of TOP1cc entrapment (derived from densitometric analysis of RADAR assay data normalized to the untreated sample) in U2-OS cells treated with different kinase inhibitors at different concentrations (**detailed in Supplementary Table 1**). (Z1) Dot blots for TOP1cc stabilization in U2-OS cells treated with CHK1 inhibitors SCH900776 (50, 100 and 250 nM) and UCN-01 (10, 25 and 50 nM) for 2 h, followed by RADAR assay. Densitometric quantification is presented at the bottom of each blot. (Z2) TOP1cc immunofluorescence in U2-OS cells. U2-OS cells were treated with 250 nM SCH, 50 nM UCN-01 (2 h) or 1 µM CPT (1 h) followed by immunofluorescent detection with monoclonal anti-TOP1cc antibody. Zoomed image is presented here. For the full field view, refer to Figure S1E. (Z3-Z5) FRAP kinetics of TOP1 in U2-OS cells. U2-OS cells were treated with 250 nM SCH for 2 h, followed by quantitative measurement of TOP1 kinetics employing FRAP. Scale bars: 10 µm. *** *p<*0.001.

We have previously reported that TOP1cc removal by WRN results in CHK1-mediated activation of the NF-κB pathway^7^ in response to nanomolar concentrations of CPT. This, in conjunction with multiple other reports have established a downstream effector function of the CHK1 kinase in cellular response to TOP1 poisoning. However, stabilization of TOP1ccs in response to ATR/CHK1 inhibition constituted a novel observation, which prompted us to investigate further a possible upstream, and more direct role played by the ATR/CHK1 pathway in regulation of cellular TOP1cc levels.

### CHK1 regulates endogenous levels of TOP1ccs in a replication and transcription-independent manner

In order to validate the findings of the kinase screen, we inhibited CHK1 catalytic activity in U2-OS cells using two different ATP-competitive small-molecule CHK1 inhibitors [SCH900776 (SCH) and UCN-01], followed by quantification of cellular TOP1cc levels using RADAR assay. Pharmacological inhibition of CHK1 with either SCH or UCN-01 resulted in extensive stabilization of TOP1ccs (**Figure 1Z1, S1B**), without affecting overall cellular TOP1 levels (**Figure S1C**). Considering the superior specificity of SCH for CHK1^27^, further investigation was carried out with SCH (henceforth referred to as CHK1*i*). CHK1*i*-mediated robust stabilization of TOP1ccs was also observed in four different cell lines of pancreatic, lung, colorectal and breast cancer origins (PANC-1, A549, HCT116, and MDA-MB-231, respectively) (**Figure S1D**). We also confirmed CHK1*i*-mediated stabilization of TOP1ccs *in situ* using immunofluorescence microscopy employing a monoclonal antibody which specifically detects TOP1ccs (**Figure 1Z2, S1E**). Finally, in order to quantitatively evaluate cellular TOP1 dynamics, we performed FRAP (fluorescence recovery after photobleaching) of EGFP-TOP1 in the presence or absence of CHK1*i*. To this end, we developed a subline of U2-OS cells with stable ectopic expression of EGFP-TOP1 (U2-OS^EGFP-TOP1^). We subjected U2-OS^EGFP-TOP1^ cells to CHK1*i* treatment and performed quantitative kinetic measurements using FRAP assay employing a laser scanning confocal microscope. Under untreated conditions, fluorescence recovery curves revealed that approximately 12-15% of nuclear TOP1 is immobile, while 85-88% of the TOP1 population is highly mobile. In contrast, CHK1*i* treatment (250 nM, 4 h) elevated the immobile population of TOP1 to ∼25-28 % in U2-OS^EGFP-TOP1^ cells, suggesting a significant change in TOP1 dynamics in response to CHK1 inhibition (**Figure 1Z3-Z4)**. Quantification of the t_1/2_ of fluorescence recovery revealed a similar phenomenon, with a t_1/2_ of ∼20 s in untreated cells as compared to ∼35 s in CHK1*i*-treated cells (**Figure 1Z5)**.

TOP1 is intimately associated with replication as well as transcription, and CHK1 is reported to play vital regulatory roles in both processes^28^. Hence, it is essential to rule out the contribution of CHK1*i*-mediated replication or transcription perturbation towards TOP1cc stabilization. First, the effect of replication on CHK1*i*-induced TOP1cc stabilization was assessed in the presence of EdU. CHK1*i* treatment led to enhanced accumulation of TOP1cc in both EdU positive (replicating) and EdU negative (non-replicating) cells (**Figure S2A, B**). Moreover, aphidicolin (a DNA polymerase α inhibitor), PHA-767491 (CDC7 inhibitor; blocks origin firing), or serum starvation reduced EdU incorporation significantly without appreciable interference with CHK1*i*-mediated TOP1cc stabilization (**Figure S2C-E, S3A-B**). Interestingly, treatment of isolated mouse splenocytes (which show negligible replication under unstimulated condition) with CHK1*i* led to extensive stabilization of TOP1ccs in a time dependent manner (**Figure S3C**). Next, we proceeded to assess the contribution of transcriptional perturbations on CHK1*i*-induced TOP1cc stabilization. We treated U2-OS cells with either 5,6-dichloro-1-beta-D-ribofuranosylbenzimidazole (DRB) or α-amanitin prior to CHK1*i* treatment. Pre-treatment of cells with either transcription inhibitor led to significant reduction in 5-Ethynyl uridine (EU) incorporation (**Figure S4A, B**) while CHK1*i*-mediated TOP1cc stabilization was unaffected (**Figure S4C**). Taken together, our results suggest that CHK1*i*-mediated TOP1cc stabilization has *per se* no discernible dependence on replication/transcription.

Cellular TOP1cc levels are tightly regulated through a fine balance between the rates of TOP1cc formation (a characteristic feature of the TOP1 catalytic cycle, determined by the relative rates of cleavage and religation) and removal (mediated by an elaborate array of post translational modifications (PTMs) and their downstream effector signals). Perturbations in either (a) the TOP1 catalytic cycle or (b) the rate of TOP1cc clearance may result in elevated steady state levels of TOP1ccs inside cells. Our previous observations prompted us to dissect the relative contribution of CHK1 in these two processes in the context of CHK1*i*-induced TOP1cc stabilization. To this end, we performed a series of experiments. CPT-mediated stabilization of TOP1ccs is known to induce global TOP1 downregulation in cells^12,29^. Hence, we first probed if CHK1*i* affects CPT-induced global TOP1 downregulation or TOP1cc clearance. We treated cells with CPT in the absence or presence of CHK1*i*. Pre-treatment with CHK1*i* (1 h) failed to prevent CPT-induced global downregulation of TOP1 (**Figure S5A**) or clearance of TOP1ccs (**Figure S5B**). Interestingly, CHK1*i* treatment did not induce global TOP1 downregulation in spite of TOP1cc stabilization. Finally, we employed DUST assay^13^ to probe post-translational modifications on TOP1ccs in CPT-treated cells with or without pre-treatment with CHK1*i*. In agreement with prevalent reports, CPT treatment triggered multiple PTMs on TOP1ccs, namely SUMOylation, ubiquitination as well as PARylation^13^ (**Figure S5C**). In line with our previous findings, CPT-triggered degradative signals on TOP1ccs were unperturbed by CHK1*i* pre-treatment (**Figure S5C**), suggesting that CHK1*i*-mediated TOP1cc stabilization is not due to perturbation of cellular TOP1cc clearance machinery.

### CHK1 interacts with and phosphorylates TOP1

In order to uncover the mechanism underlying the above observations, we investigated the interaction between CHK1 and TOP1 with a series of experiments. Initially, we employed alphafold multimer (AFM) for a predictive exploration of TOP1-CHK1 interaction. In order to eliminate interferences from the N-terminal disordered region of TOP1, we used amino acids 201-765 for analysis. Analysis of AFM outputs revealed a moderately confident prediction of TOP1-CHK1 interaction, with two models satisfying parametric benchmarks of model quality (**Figure 2A-D**, **S6A-F**). Additionally, we separately probed the interaction between the kinase (amino acids 1-265) and KA1 (amino acids 391-476) domains of CHK1 with TOP1. Imperatively, only the CHK1 kinase domain, but not the KA1 domain (which is associated with the regulation of CHK1 kinase domain) was predicted to interact with TOP1 (**Figure S6G-R)**. We followed up these theoretical predictions with validation of the interaction between TOP1 and CHK1 at cellular level. We transiently overexpressed EGFP-TOP1 in U2-OS cells and performed immunoprecipitation with anti-GFP antibody. CHK1 was readily co-immunoprecipitated with EGFP-TOP1 (**Figure 2E**). This was further confirmed using proximity ligation assay (**Figure 2F, G**). TOP1-CHK1 interaction was unabated in the presence of CHK1*i* in both cases, which is in line with previously reported non-interference of ATP-competitive CHK1 inhibitors with CHK1-substrate interactions^30^. In order to assess whether this interaction leads to phosphorylation of TOP1, purified full length TOP1 was incubated with CHK1 in the presence of γ^32^P-ATP. CHK1 phosphorylated TOP1 *in vitro,* confirming that TOP1 is indeed a substrate for CHK1 (**Figure 2H**).

**Figure 2:**
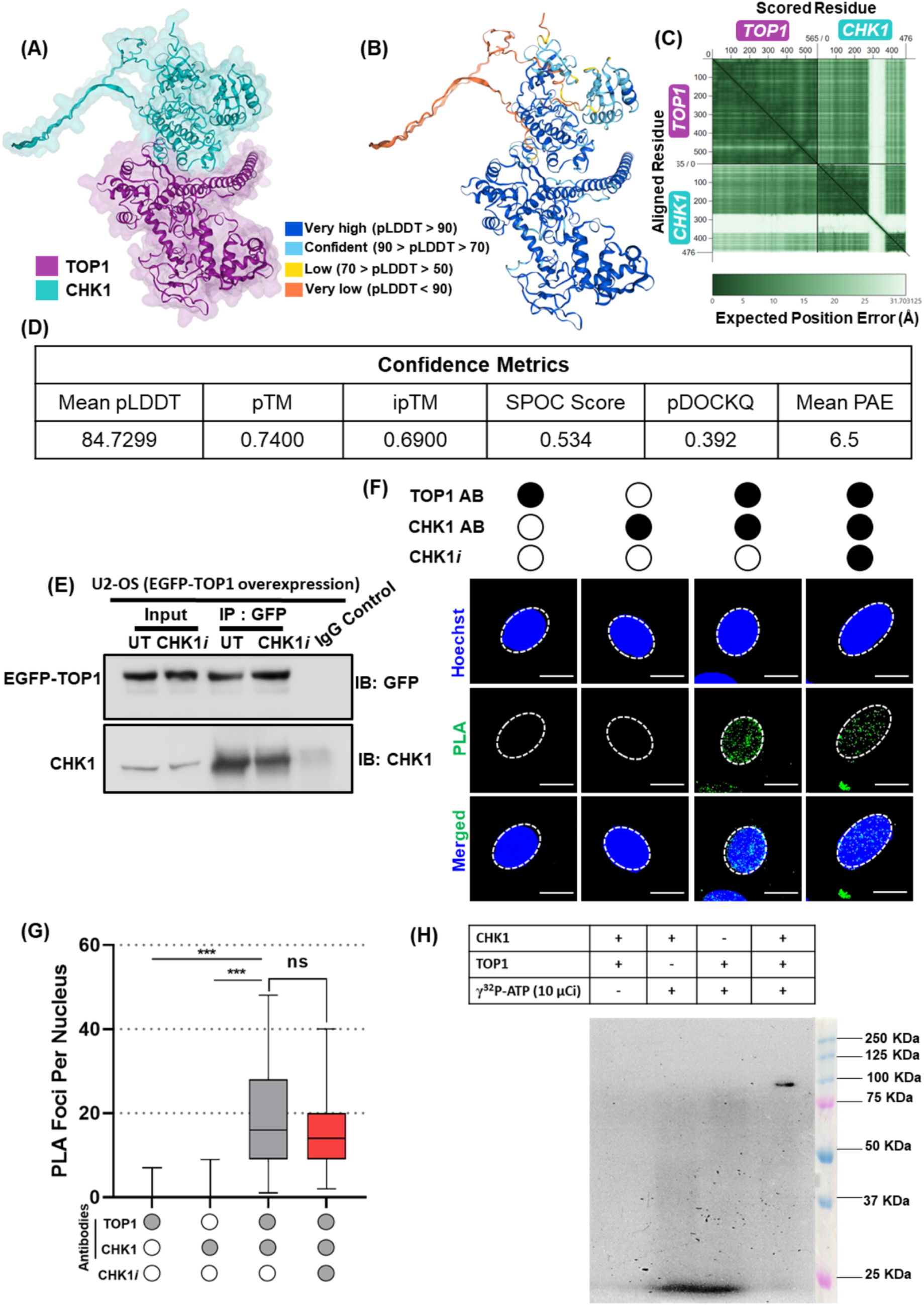
Validation of CHK1-TOP1 interaction *in situ* and *in vitro* phosphorylation of TOP1 by CHK1. (A,B) Alphafold multimer prediction of TOP1-CHK1 interaction. The top ranked model has been reproduced (A – Subunit view, B – pLDDT view). (C) PAE plot of the top ranked model. (D) Confidence metrics of the predicted TOP1-CHK1 interaction. While pLDDT, pTM and iPTM were calculated based on the top ranked model, SPOC, pDOCKQ and mean PAE were calculated based on the three top ranked models. Tolerance cutoff of each parameter is as follows: pLDDT<75, pTM>0.7, ipTM>0.55, SPOC>0.3, pDOCKQ>0.23, PAE<15. (E) Co-immunoprecipitation of TOP1 and CHK1 in U2-OS cells. U2-OS cells were transiently transfected with EGFP-TOP1. TOP1 was immunoprecipitated from transfected cells (untreated, or treated with 250 nM CHK1*i* for 2 h) with anti-GFP antibody. (F, G) Proximity ligation assay of TOP1 and CHK1 in U2-OS cells with or without treatment with 250 nM CHK1*i* for 2 h. (H) *In vitro* kinase assay with TOP1 and CHK1. Abbreviations: pLDDT – Predicted Local Distance Difference Test, pTM – Predicted Template Modelling score, ipTM – Interface Template Modelling score, SPOC – Structure Prediction and Omics Classifier, PAE – Predicted Aligned Error. Scale bars: 10 µm. *** *p<*0.001.

### CHK1 phosphorylates Serine 320 of TOP1, which regulates TOP1 dynamics

Based on our observation that CHK1 interacts with and phosphorylates TOP1, we proceeded to predict putative CHK1-targeted consensus sequences on TOP1. We employed the experimentally validated CHK1-specific serine/threonine-containing motifs derived by Blasius and coworkers^31^ to perform a sequence motif search on TOP1 protein using the Prosite server (https://prosite.expasy.org/). *In silico* analysis revealed two putative CHK1 target sites on TOP1: T154 and S320 (**Figure 3A**), neither of which were previously reported to be phosphorylated by any kinase. In another analysis, when a minimal CHK1 target motif was used as query, multiple additional consensus sites were identified on TOP1 (**Supplementary Table 2**). In order to investigate which amino acids are phosphorylated on TOP1 under physiological conditions, we designed a tandem Mass Spectrometry (MS)-based assay (**Figure 3B**). To specifically detect sites phosphorylated on catalytically engaged TOP1, TOP1ccs were purified using DUST assay workflow (with untreated, CHK1*i* or CPT treated U2-OS cells). This was followed by tandem-MS analysis of excised bands from stained SDS PAGE gels. MS analysis led to the detection of multiple phosphorylation sites (Y231, S250, S320, S394, T446, Y461, Y480, S534, T570, Y538, Y706) in untreated or treated samples (CHK1*i*, CPT) at different time points (**Supplementary Table 3**; **Figure 3C**). Among four predicted CHK1-mediated TOP1 phosphorylation sites (T154, S320, S394 and T570), T154 was not detected in our MS-analysis. In order to understand the physiological relevance of these phosphosites, we generated three “phospho-resistant” mutants of TOP1 (EGFP-TOP1^S320A^, EGFP-TOP1^S394A^ and EGFP-TOP1^T570A^) followed by evaluation of their dynamics on genomic DNA employing RADAR assay. RADAR assay with anti-GFP antibody revealed similar changes in EGFP-TOP1^WT^ dynamics compared to endogenous TOP1 in response to CHK1*i* (**Figure 3D** *vs* **Figure 1Z1**). Intriguingly, EGFP-TOP1^S320A^ was constitutively trapped on the genome (**Figure 3D**) in untreated cells. CHK1*i* treatment did not result in further enhancement of stabilization of EGFP-TOP1^S320A^, which was expected, given the copious stabilization of EGFP-TOP1^S320A^ under untreated conditions. In contrast, the kinetics of EGFP-TOP1^S394A^ and EGFP-TOP1^T570A^ on the genome were essentially identical to that of EGFP-TOP1^WT^ under untreated or CHK1*i* treated conditions (**Figure 3D**). Thus, our results suggested that while phosphorylation at S320 plays a critical role in regulating TOP1 dynamics, T570 or S394-phosphorylation is unlikely to be involved in determining cellular levels of TOP1ccs under ambient conditions. Notably, both EGFP-TOP1^WT^ and EGFP-TOP1^S320A^ were downregulated in response to CPT (**Figure 3E**), indicating these proteins are functionally active at cellular levels. Moreover, our homology search revealed that TOP1-S320 is highly conserved in species ranging from yeast to vertebrates (**Figure 3F**). Structurally, S320 is strategically located in a connecting loop between the α5 and α6 helices, which constitute the “nose-cone” resident within the “Cap” domain of TOP1 (**Figure 3G**). The nose-cone helices have been shown to make multiple contacts with the substrate downstream of the scissile site, and play an indispensable role in the TOP1 catalytic cycle^32–34^. Together, our results demonstrated a critical role of CHK1-mediated phosphorylation of TOP1 at S320 in regulating TOP1 dynamics at physiological level.

**Figure 3:**
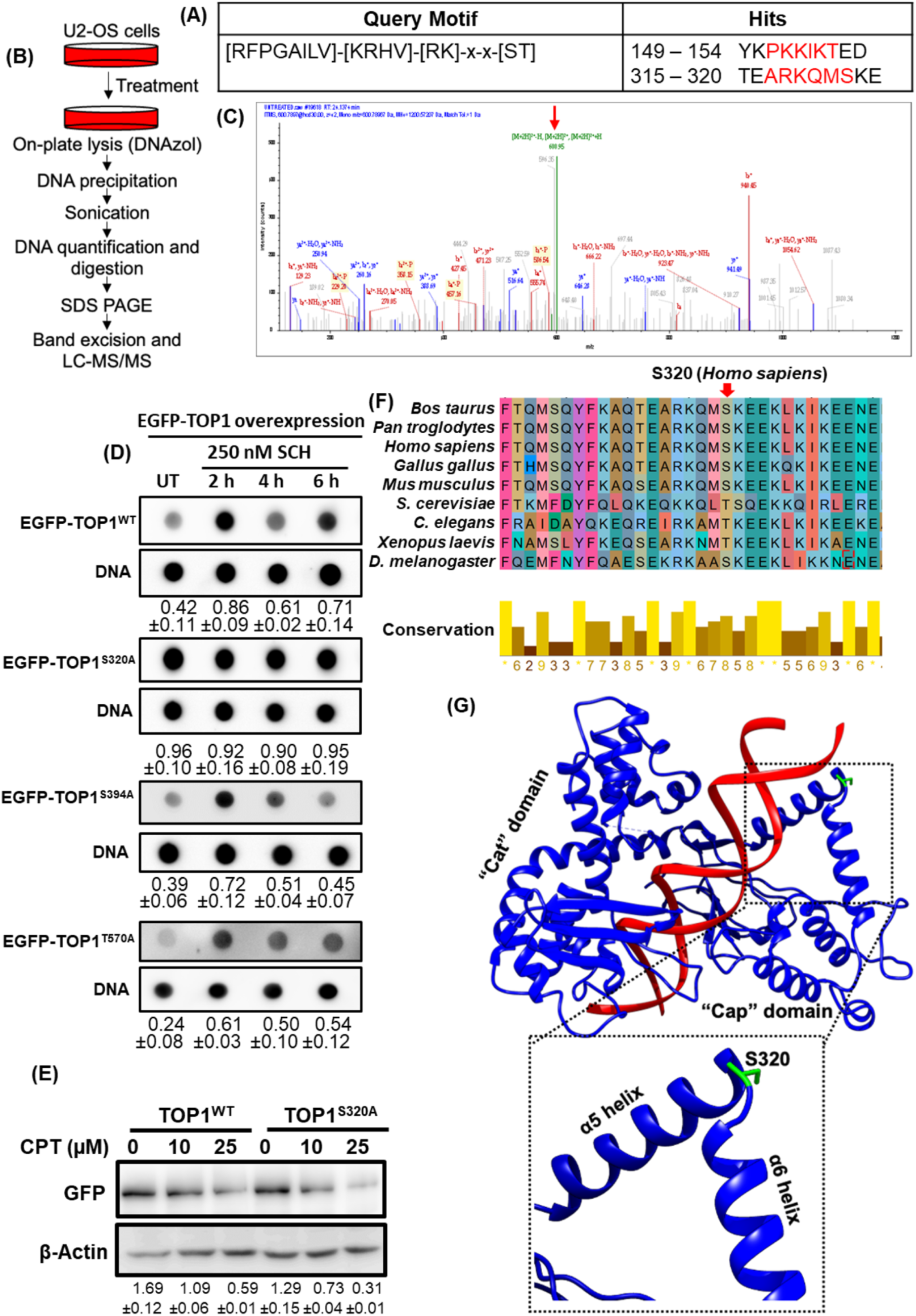
Prediction of CHK1 target sites on TOP1 and mass spectrometric detection of TOP1 post translational modifications. (A) Results of CHK1 target motif search on TOP1. (B) Workflow for the mass spectrometric detection of post translational modifications on catalytically engaged TOP1. (C) Representative spectra of a peptide encompassing the phosphorylated Serine 320 residue on TOP1. (D) RADAR assay in U2-OS cells transfected with EGFP-TOP1^WT^, EGFP-TOP1^S320A^, EGFP-TOP1^S394A^ or EGFP-TOP1^T570A^. U2-OS cells were transfected with EGFP-TOP1^WT^, EGFP-TOP1^S320A^, EGFP-TOP1^S394A^ or EGFP-TOP1^T570A^ followed by treatment with 250 nM CHK1*i* (for 2, 4 or 6 h), 24 h post transfection. Dot blots were probed with anti-GFP antibody. (E) Global downregulation of EGFP-TOP1^WT^ and EGFP-TOP1^S320A^. U2-OS cells were transiently transfected with EGFP-TOP1^WT^ and EGFP-TOP1^S320A^, followed by treatment with indicated concentrations of CPT for 2 h. Downregulation of wild type or S320A mutant was followed by probing for GFP. (F) Multiple sequence alignment of TOP1 from different eukaryotic species. (G) Structural context of Serine 320 in the TOP1 protein.

### CHK1 regulates TOP1 catalytic activity

In order to further confirm whether CHK1 regulates TOP1 catalytic activity, we performed plasmid relaxation assays with crude nuclear extracts prepared from untreated and CHK1*i*-treated U2-OS cells. CHK1*i* treatment resulted in significant depression in cellular TOP1 catalytic activity (**Figure 4A, B**). Moreover, CHK1*i* showed no direct catalytic inhibition of purified human TOP1 (**Figure 4C-D**), ruling out any non-specific direct inhibitory effect of CHK1*i* on TOP1. Further, EGFP-TOP1^WT^ was immunoprecipitated from untreated and CHK1*i*-treated cells (expressing ectopic EGFP-TOP1^WT^). Western blot analysis was performed to ensure equal protein expression levels and immunoprecipitation efficiencies (**Figure 4E**). Time kinetics of plasmid relaxation assays revealed that treatment of cells with CHK1*i* significantly compromises EGFP-TOP1^WT^ catalytic activity (**Figure 4F, G**). In order to specifically dissect the role played by S320 phosphorylation in TOP1 catalysis, EGFP-TOP1^WT^ as well as EGFP-TOP1^S320A^ were immunoprecipitated (**Figure 4H**) and their relative catalytic activities were assessed employing plasmid relaxation assay. Intriguingly, EGFP-TOP1^S320A^ had significantly compromised catalytic activity compared to EGFP-TOP1^WT^ (**Figure 4I, J**), essentially producing a phenocopy of CHK1*i*-mediated depression of TOP1 catalysis. Our results establish that basal cellular CHK1 activity plays a critical role in directly regulating TOP1 catalytic activity through phosphorylation of S320.

**Figure 4:**
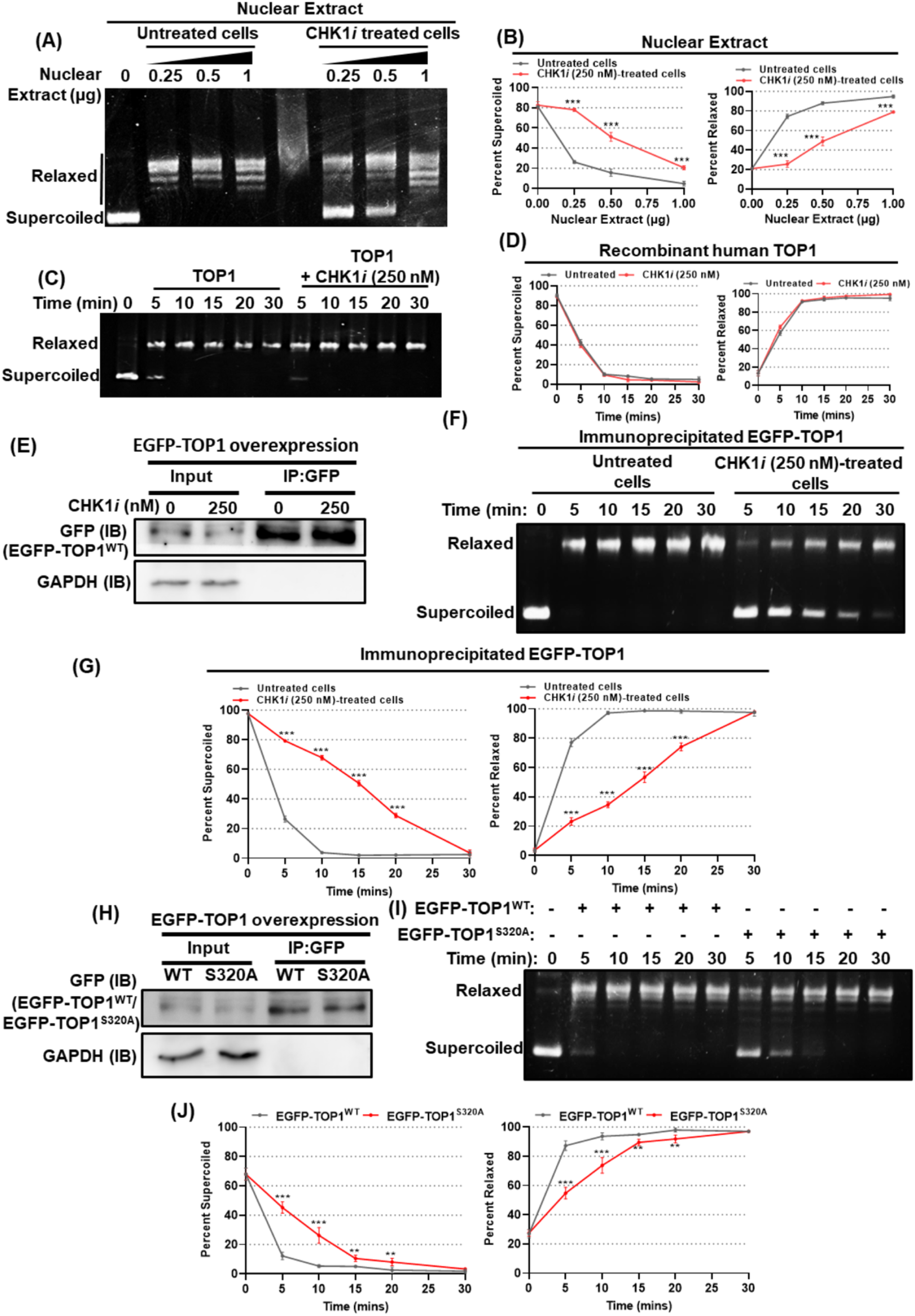
CHK1 regulates TOP1 catalytic activity in cells. (A, B) Plasmid relaxation assay with crude nuclear extracts prepared from U2-OS cells (untreated, or treated with 250 nM CHK1*i* for 2 h). (C, D) Plasmid relaxation assay with recombinant human TOP1 in the absence or presence of 250 nM CHK1*i*. (E-G) Plasmid relaxation assay with immunoprecipitated EGFP-TOP1^WT^. U2-OS cells were transiently transfected with EGFP-TOP1^WT^. 48 h post transfection, cells were treated with 250 nM CHK1*i* for 2 h, followed by immunoprecipitation of EGFP-TOP1 from untreated with CHK1*i*-treated cells. Immunoblot was performed to confirm the immunoprecipitation efficiency. Plasmid relaxation assays were performed with the isolated immunocomplexes. (H-J) Plasmid relaxation assay with EGFP-TOP1^WT^ and EGFP-TOP1^S320A^. U2-OS cells were transiently transfected with EGFP-TOP1^WT^ and EGFP-TOP1^S320A^, followed by immunoprecipitation 48 h post transfection. Immunoprecipitation efficiency was confirmed using immunoblotting, and plasmid relaxation assay was performed with the isolated immunocomplexes. *** *p<*0.001.

### CHK1-mediated TOP1 regulation limits replication-associated genomic instability

TOP1 poisoning-associated TOP1cc stabilization has been widely established to induce genomic instability^1,35^. Since TOP1^S320A^ is catalytically compromised and constitutively trapped on genomic DNA, we envisaged that CHK1-mediated TOP1 phosphorylation (S320) might play a critical role in limiting TOP1cc-associated genomic instability. In order to examine this, we evaluated the basal DNA damage levels in U2-OS sublines with stable ectopic expression of EGFP-TOP1^WT^ and EGFP-TOP1^S320A^ (U2-OS^EGFP-TOP1WT^ and U2-OS^EGFP-TOP1S320A^, respectively). U2-OS^EGFP-TOP1S320A^ cells had significantly higher levels of γ-H2AX as well as 53BP1 foci, indicating higher basal DNA damage levels (**Figure 5A-C, S7A**). In order to delineate the role of replication in TOP1^S320A^-associated DNA damage, we inhibited DNA replication in U2-OS^EGFP-TOP1S320A^ cells (with aphidicolin or CDC7 inhibitor, respectively), followed by evaluation of DNA damage levels. Replication inhibition significantly alleviated DNA damage levels in U2-OS^EGFP-TOP1S320A^ cells (**Figure 5D, E**), demonstrating that TOP1^S320A^-mediated DNA damage is partially associated with replication. Additionally, EdU incorporation assay revealed a significant impairment in EdU incorporation in U2-OS^EGFP-TOP1S320A^ cells *vis-à-vis* U2-OS^EGFP-TOP1WT^ cells (**Figure S7B, C**). In order to assess direct impact of TOP1-S320A mutation on replication, single molecule DNA fiber analysis was performed in U2-OS cells transiently transfected with EGFP-TOP1^WT^ and EGFP-TOP1^S320A^. EGFP-TOP1^S320A^ expression resulted in severe impairment of replication fork progression, evident from shortening of fiber track lengths (**Figure 5F-H**). However, no significant changes in fork symmetry was observed **Figure 5I**). Moreover, in concordance with these results, U2-OS^EGFP-TOP1S320A^ cells had a slower growth rate compared to U2-OS^EGFP-TOP1WT^ cells (**Figure 5J**). Finally, we prepared metaphase spreads from U2-OS^EGFP-TOP1WT^ and U2-OS^EGFP-TOP1S320A^ cells in order to investigate gross chromosomal abnormalities (if any). Quantitative scoring of dicentrics, acentric fragments and chromatid breaks revealed that U2-OS^EGFP-TOP1S320A^ cells suffer severely from higher levels of chromosomal abnormalities, suggestive of large-scale genomic instability (**Figure 5K, L**). Taken together, our data suggests a critical role played by S320 phosphorylation is limiting TOP1 associated genomic instability.

**Figure 5:**
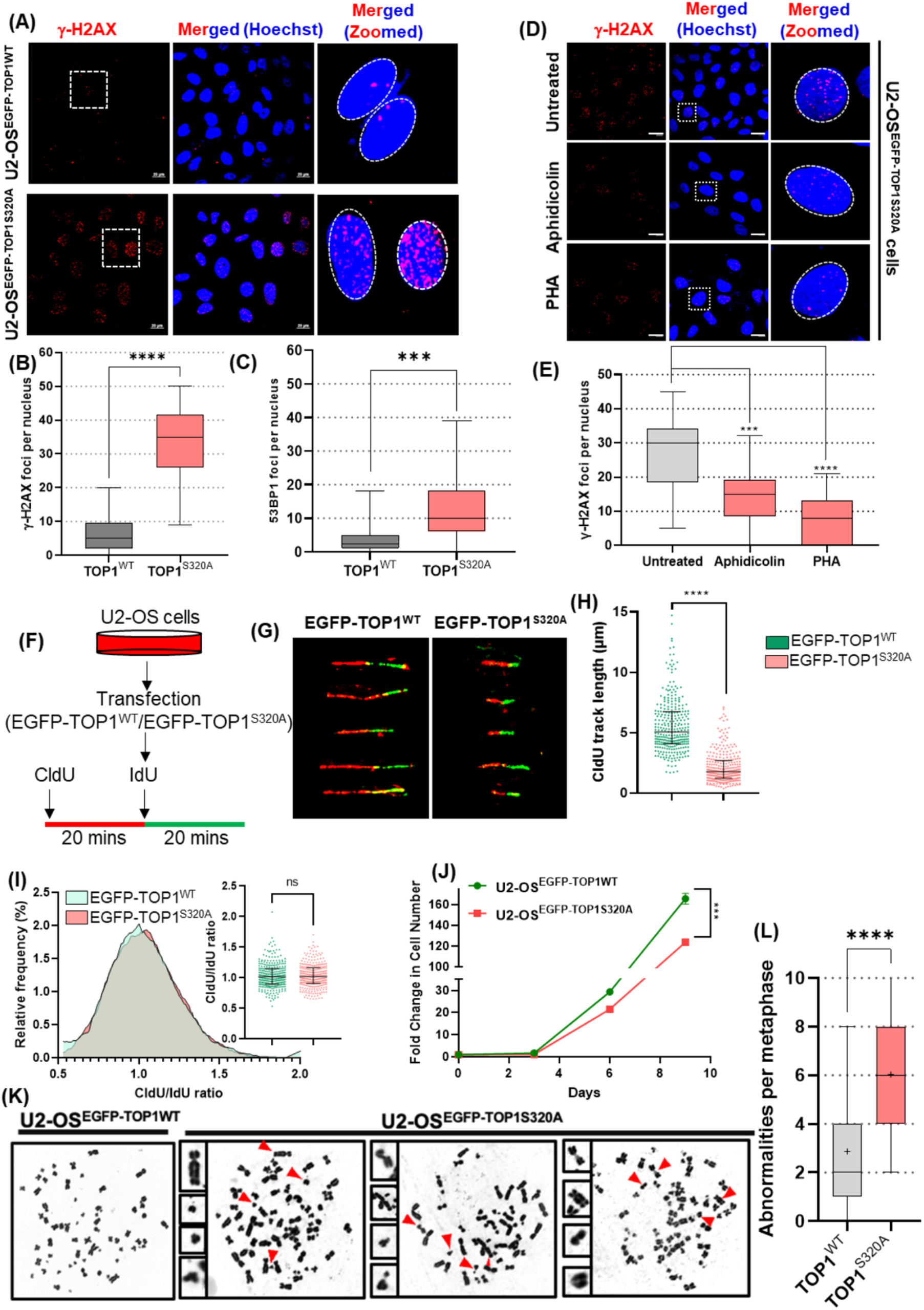
The effects of TOP1-S320 mutation on DNA damage, growth kinetics and genomic instability. (A-C) Estimation of DSB (of γ-H2AX and 53BP1) levels in U2-OS^EGFP-TOP1WT^ and U2-OS^EGFP-TOP1S320A^ stable cells. (D-E) γ-H2AX levels in U2-OS^EGFP-TOP1S320A^ cells treated with replication inhibitors Aphidicolin or CDC7*i* (PHA). Cells were treated with 200 nM aphidicolin for 16 h or 5 µM PHA for 6 h before immunofluorescent detection of γ-H2AX foci. (F-G) DNA fiber assay with U2-OS cells transiently transfected with EGFP-TOP1^WT^ and EGFP-TOP1^S320A^. Five representative DNA fibers have been shown per sample. (H) Quantitative analysis of CldU track lengths. (I) Histogram of CldU/IdU track length ratios representing fork symmetry. Statistical analysis of the same data has been shown in inset. (J) Growth kinetics of U2-OS^EGFP-TOP1WT^ and U2-OS^EGFP-TOP1S320A^ stable cells. U2-OS^EGFP-TOP1WT^ and U2-OS^EGFP-TOP1S320A^ cells were seeded in 60 mm dishes and cell number was quantified 3, 6 or 9 days post seeding. (K-L) Representative metaphase spreads prepared from U2-OS^EGFP-TOP1WT^ and U2-OS^EGFP-TOP1S320A^ cells (and statistical analyses thereof). Scale bars: 20 µm. *** *p<*0.001; **** *p<*0.0001.

### CHK1 supresses TOP1-associated R-loop accumulation and transcription-replication collisions

Stabilization of TOP1ccs by TOP1 poisons has been widely shown to stabilize R-loops and induce R-loop-associated genomic instability^3,35,36^. In order to investigate whether TOP1ccs arising from compromised catalytic activity of TOP1^S320A^ translate into transcription-dependent DSBs, we measured γ-H2AX levels in U2-OS^EGFP-TOP1S320A^ cells in the presence or absence of the transcription inhibitor DRB. Treatment with 100 µM DRB suppressed DSBs in U2-OS^EGFP-TOP1S320A^ cells in a time dependent manner (**Figure 6A, B**), suggesting that constitutive stabilization of TOP1(S320A)ccs indeed results in transcription-dependent DNA damage. Next, we measured global transcription levels in U2-OS^EGFP-TOP1WT^ versus U2-OS^EGFP-TOP1S320A^ cells through EU incorporation assay. U2-OS^EGFP-TOP1S320A^ cells showed a mild (but significant) depression in global transcription levels (**Figure S7C, D**). We further measured the effects of TOP1(S320A)ccs on global R-loop levels. Microscopic detection of R-loops through S9.6 monoclonal antibody revealed that U2-OS^EGFP-TOP1S320A^ cells harbour significantly higher levels of R-loops compared to U2-OS^EGFP-TOP1WT^ cells (**Figure 6C-D**). S9.6 specific signals were drastically reduced, when cells were treated with RNaseH (which degrades RNA-DNA hybrids), confirming the specificity of S9.6 antibody for R-loop structures. Dot blot-based R-loop analysis was also performed with isolated pooled DNA from cells (**Figure 6E**). We observed similar higher levels of R-loops in U2-OS^EGFP-TOP1S320A^ cells compared to U2-OS^EGFP-TOP1WT^ cells (**Figure 6E, F**). CPT-mediated TOP1cc stabilization has been shown to trigger transcription-replication collisions (TRCs)^37^. Hence, we further analysed the levels of TRCs in U2-OS^EGFP-TOP1WT^ and U2-OS^EGFP-TOP1S320A^ cells employing PLA between replisome-associated protein PCNA and elongating RNA pol II (phosphoserine 2). PLA results revealed significant prevalence of TRCs in U2-OS^EGFP-TOP1S320A^ cells, in contrast to U2-OS^EGFP-TOP1WT^ cells (**Figure 6G, H**). Hence, our results indicate that CHK1 suppresses transcription-associated genomic instability (R-loop and DSBs) through phosphorylation-mediated regulation of TOP1 activity at physiological level.

**Figure 6:**
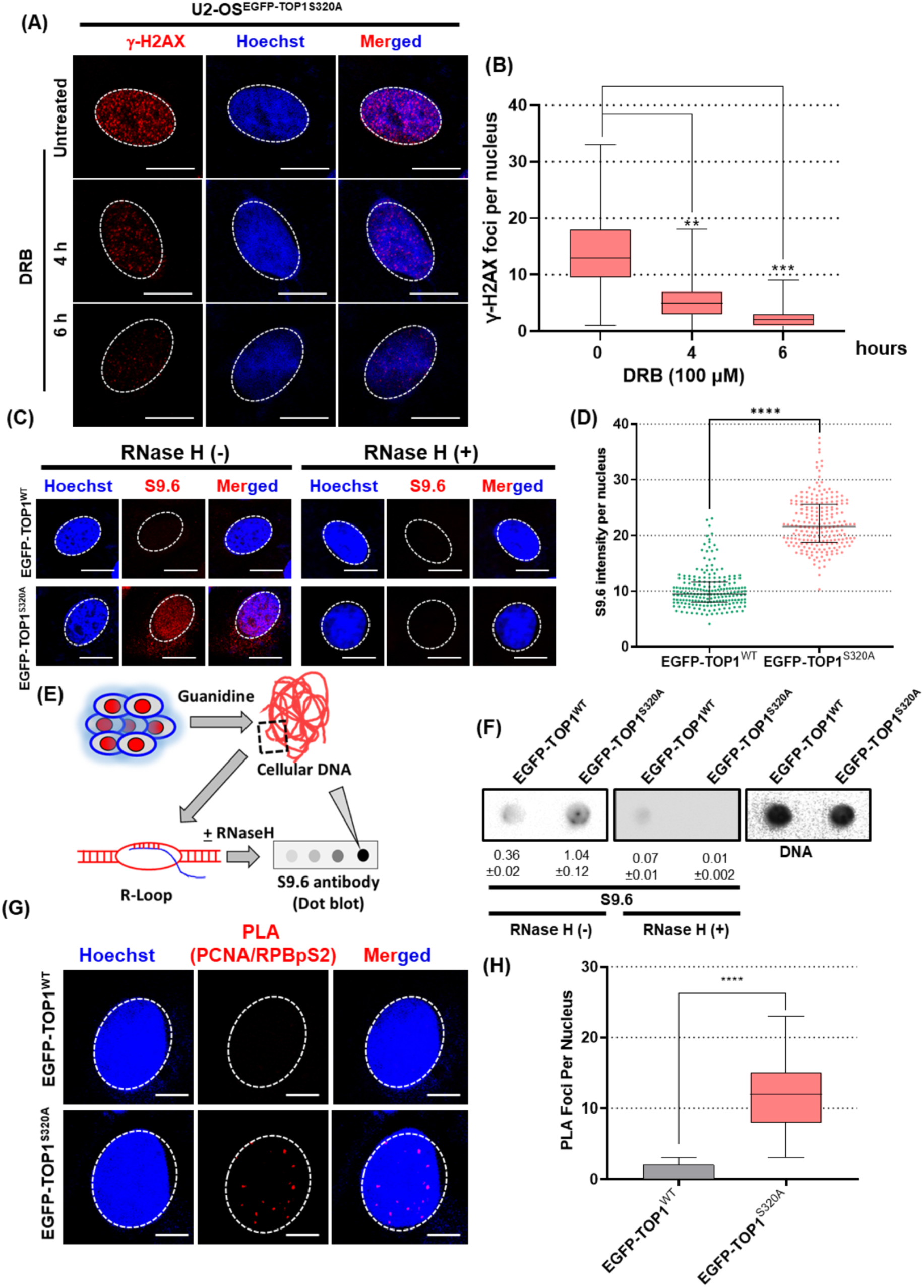
The effects of TOP1-S320 mutation on transcription-associated DNA damage, R-loop accumulation and transcription-replication collisions in cells. (A, B) DNA damage levels in U2-OS^EGFP-TOP1S320A^ in the presence of the transcription inhibitor DRB. U2-OS^EGFP-TOP1S320A^ cells were treated with 100 µM DRB for 4 h or 6 h followed by immunofluorescent detection of γ-H2AX foci. (C, D) RNA/DNA hybrid (R-loop) levels in U2-OS^EGFP-TOP1WT^ and U2-OS^EGFP-TOP1S320A^ stable cells. R-loops were immunodetected with the S9.6 monoclonal antibody. Signal specificity was determined using RNase H. (E, F) Dot blot of R-loop levels in U2-OS^EGFP-TOP1WT^ and U2-OS^EGFP-TOP1S320A^ stable cells. (G, H) Proximity ligation assay (PLA) for detection of transcription-replication collisions (TRCs) in U2-OS^EGFP-TOP1WT^ and U2-OS^EGFP-TOP1S320A^ stable cells. PLA was performed with PCNA and RNA pol II large subunit phospho-serine 2 antibodies. Scale bars: 10 µm. *** *p<*0.001; **** *p<*0.0001.

### CHK1*i* preferentially downregulates the expression of long genes through stabilization of TOP1ccs and R-loops at gene bodies

CPT-mediated stabilization of TOP1ccs has been reported to result in preferential downregulation of long and highly expressed genes^38,39^. We have demonstrated that the phospho-resistant TOP1^S320A^ mutant is constitutively trapped on the genome, and induces transcription-associated DNA damage as well as accumulation of deleterious R-loops. These findings prompted us to investigate the transcriptional effects of CHK1*i vis-à-vis* short and long genes. To this end, we performed transcriptomic analysis (∼20000 genes) of cells treated with CHK1*i* for 2 or 6 h. CPT (1 µM, 2 h) treatment was included as a positive control. Interestingly, close inspection of lengths of genes which were preferentially downregulated versus those whose expression was unaltered revealed that CHK1*i* treatment recapitulates CPT-mediated preferential downregulation of long and highly expressed genes **(Figure 7A**). Further, we sought to investigate the spatial distribution of TOP1^S320A^ in the transcriptional context of genes. To this end, we employed a previously reported panel of probes^3^ pertaining to promoter proximal regions (PPR) and gene bodies of a few well characterized genes (three and six genes, respectively). As an additional measure, we also included probes against MYO3A which was the longest gene (2.1 megabases) found to be downregulated in response to CHK1*i* treatment. We ectopically expressed EGFP-TOP1^WT^ and EGFP-TOP1^S320A^ in U2-OS cells and performed chromatin immunoprecipitation from uncrosslinked nuclei (**Figure 7B**)^40^. EGFP-TOP1^S320A^ had significantly higher chromatin occupancy than EGFP-TOP1^WT^ at gene bodies (**Figure 7C**). However, such an effect was not seen at promoter proximal regions. Interestingly, this is in line with previously reported preferential enrichment of catalytically active TOP1 on gene bodies over promoter proximal regions^40^, which suggests that while EGFP-TOP1^S320A^ remains catalytically active, its slower dynamics on the chromatin contributes to enhanced levels of TOP1(S320A)ccs on gene bodies. We further envisaged that R-loops generated due to elevated TOP1(S320A)ccs should spatially and physically coincide with TOP1(S320A)ccs. To test this hypothesis, we performed DNA/RNA hybrid immunoprecipitation (DRIP) with U2-OS cells ectopically expressing EGFP-TOP1^WT^ and EGFP-TOP1^S320A^, and performed quantitative real time PCR with the above probes. In agreement with our reasoning, there was significantly higher R-loop enrichment in EGFP-TOP1^S320A^ samples. Further, with the only exception of geminin, R-loops were spatially colocalized with TOP1ccs in gene bodies (**Figure 7D**). Taken together, these results shed important light on the genome wide transcriptional effects of CHK1 inhibition and establish a transcriptional context of TOP1cc stabilization and R-loop accumulation in response to the same.

**Figure 7:**
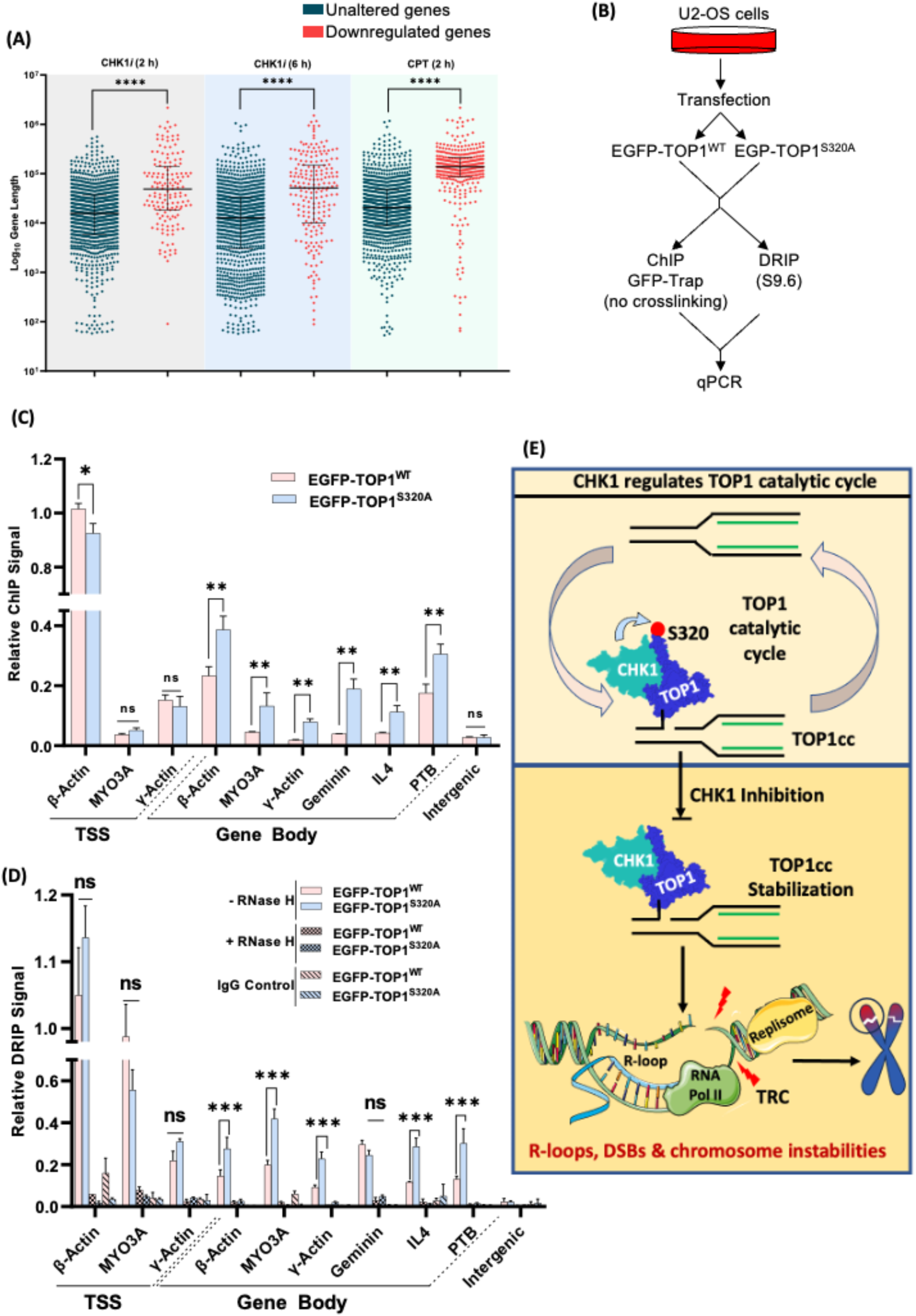
Global transcriptional effects of CHK1 inhibition *vis-à-vis* gene length and transcriptional context of genes. (A) Analysis of gene lengths post transcriptomic analysis of U2-OS cells treated with CHK1*i* (250 nM) or CPT (1 µM). (B) Workflow for ChIP and DRIP assays. (C) Relative enrichment of TOP1ccs in transcription start sites (TSS) and gene bodies arising from EGFP-TOP1^WT^ and EGFP-TOP1^S320A^. U2-OS cells were transiently transfected with EGFP-TOP1^WT^ and EGFP-TOP1^S320A^, followed by chromatin immunoprecipitation with anti-GFP antibody from un-crosslinked nuclei. (D) Relative enrichment of R-loops at TSS and gene bodies. U2-OS cells were transiently transfected with EGFP-TOP1^WT^ and EGFP-TOP1^S320A^, followed by immunoprecipitation with S9.6 monoclonal antibody. Specificity of signal was determined using RNase H sensitivity. (E) A schematic model for the maintenance of genomic stability by CHK1 through regulation of TOP1 catalytic cycle. Under unperturbed physiological conditions (top panel), CHK1-mediated phosphorylation at Serine 320 of TOP1 results in optimal execution of the TOP1 catalytic cycle, resulting in trace levels of TOP1ccs inside cells. In the presence of CHK1*i* (bottom panel), disruption of TOP1 catalytic regulation leads to widespread TOP1cc stabilization. The downstream consequences of CHK1*i*-associated TOP1 stabilization include replication and transcription-mediated DNA damage, transcription-replication collisions (TRCs) and widespread RNA-DNA hybrid (R-loop) stabilization, eventually leading to chromosomal abnormalities, indicative of genomic instability. *ns* non-significant; ** *p<*0.01; *** *p<*0.001; **** *p<*0.0001.

## Discussion

Maintenance of genomic integrity is of central importance to the survival of cells. TOP1 is a conserved protein which has multiple roles in the context of vital cellular processes that involve unwinding of the DNA double helix^41^. The weakest link of the TOP1 catalytic cycle is the TOP1-DNA covalent complex (TOP1cc), which, on one hand, is an obligate intermediate, and on the other hand, a source of DNA damage and genomic instability, if not properly resolved. Hence, stabilization of TOP1ccs has not only been an attractive target of anticancer therapy, but also an elegant model for studying cellular responses to DNA-protein adducts. Although elaborate mechanisms for various PTMs of TOP1ccs are known for their removal in response to TOP1 poisons^6-11^, there have been no comprehensive reports to date demonstrating how cell regulates TOP1 catalytic cycle under unperturbed physiological conditions. Deregulation of such process may have catastrophic consequences in cells, leading to genomic instability and cell death. Employing a small molecule library targeting 42 bonafide cellular kinases (**Figure 1A-Z**), the present investigation highlights a completely unknown mechanism of TOP1 regulation, wherein CHK1 mediated phosphorylation at S320 of TOP1 accelerates TOP1 catalytic cycle to minimise genome-wide accumulation of TOP1ccs in cancer cells. TOP1-S320 phosphorylation by CHK1 is found to be critical in reducing DSBs, R-loops and chromosomal instability (**Figure 7E**). Strikingly, our findings showed that TOP1ccs generated by inhibition of CHK1 is not prone to rapid removal/degradation (**Supplementary Figure S1B**), as compared to TOP1ccs generated by TOP1 poisons (**Supplementary Figure S5A,B**). Mechanistically, CPT induced TOP1ccs are recognised by PTMs that engage the proteasomal machinery for their rapid removal while CHK1*i*-mediated TOP1ccs remain stable without being recognised for longer duration (**Supplementary Figure S5C**). This may be attributable to confirmational differences in TOP1-DNA-CPT ternary complex *vs* unphosphorylated TOP1(S320)-DNA covalent complex (TOP1-S320cc) Considering the fact that TOP1ccs are lethal structures^8^, unless rapidly removed, this finding demonstrating longer stabilization of TOP1cc by CHK1*i* is intriguing and opens up a novel therapeutic strategy to target TOP1 for cancer therapy.

Previously specific *in vivo* phosphorylation of TOP1, at serine (S10, S21, S112, S394) and tyrosine (Y268, Y506) has been reported^16-19^. Of note, these TOP1 phosphorylation sites were identified from total cellular endogenous/overexpressed TOP1 (soluble mobile fraction), preventing potential identification of phosphorylation(s) on catalytically engaged TOP1, which may shed significant light on regulation of TOP1cc dynamics. To address this issue, we have used mass spectrometry to identify phosphosites exclusively on endogenous TOP1ccs at cellular level. Our result displayed a unique signature of phosphosites on catalytically engaged TOP1 (**Supplementary Table 2**), revealing significant departures from previous reports involving soluble TOP1. Multiple novel (S320, Y231 and S250) as well as known (S394, T570 and Y480) phosphosites were detected on catalytically engaged TOP1 under unperturbed conditions. In response to lower concentration of CPT (200 nM), but not higher concentration of CPT (1 μM), four additional phosphosites (Y461, S534, Y538, T706) were detected (**Supplementary Table 3**). Interestingly, presence of phosphorylation at TOP1 S320 in both untreated and CPT/CHK1*i*-treated cells, suggested its importance at physiological level. The physiological importance of S320 is highlighted by the evolutionary conservation of the site from yeast to humans. A preliminary sequence motif search on the TOP1 protein using the Prosite server (https://prosite.expasy.org/) predicted TOP1-S320 and TOP1-S394 as sites of phosphorylation by CHK1 (**Figure 3A**). A series of our experiments revealed that CHK1 interacts with TOP1 *in vitro* and *in vivo* and phosphorylates TOP1 *in vitro* (**Figure 2A-H**). Pharmacological inhibition of CHK1 or mutation of S320 in TOP1 led to extensive accumulation of TOP1ccs (**Figure 1Z1, 3E**). Furthermore CHK1*i* had no further effect on TOP1(S320A)cc formation in cells (**Figure 3E**), confirming regulation of TOP1cc dynamics through CHK1 mediated TOP-S320 phosphorylation. Intriguingly, CHK1*i* induced TOP1ccs are different than CPT-induced TOP1cc, as the later, but not CHK1*i* induced TOP1ccs, were recognised by PTM machineries for rapid removal (**Supplementary Figure S5**).

CHK1 mediated TOP1-S320 phosphorylation in unperturbed cellular conditions raises an important question: why do cells need such an intricate regulation process? In the pursuit to unravel the significance of TOP1-S320 phosphorylation by CHK1, we have carried out a series of experiments. Our data showed several important aspects: (1) TOP1 from nuclear extract of CHK1*i* treated cells or TOP1-S320A mutant is catalytically latent and sluggish in relaxing DNA (**Figure 4**), resulting in accumulation of catastrophic TOP1ccs (**Figure 1Z1, 3E**), (2) TOP1cc accumulation, due to CHK1*i* or TOP1-S320 mutation leads to severe impact on the ongoing replication and transcription process and culminates in generation of DSBs, R-loops, chromosomal breaks and fusions (**Figure 5, 6**), suggesting importance of CHK1 mediated TOP1-S320 phosphorylation in maintaining genomic stability and likely, in preventing cancers. In this regard, however, our extensive search on The Cancer Genome Atlas (TCGA) database failed to show prevalence of TOP1-S320 mutations in various cancers, which rules out direct involvement of TOP1-S320 mutation in oncogenesis. However, our study has a critical implication towards the understanding of a novel role of CHK1 in cancers. CHK1 expression is found to be low in multiple malignancies, compared to corresponding normal tissues. Multiple studies have shown that high expression of CHK1 is associated with poor prognosis, recurrence and compromised survival in breast cancer^42^, lung cancers^43^, leukemia^44^ and other cancers. Imperatively, therapy resistance in cancers is attributable to higher CHK1 levels (protein and mRNA)^45–47^. Combined treatment with CHK1 inhibitors and chemo-/radio-therapy alleviates cancer chemo/radio-resistance and abrogates tumor growth^48,49^. Hence, CHK1 inhibitors along with various chemotherapeutics are currently undergoing multiple clinical trials^25^. Based on the existing understanding of CHK1 function, anti-cancer properties of CHK1 inhibitors are associated with direct impact on replication and abrogation of cell cycle checkpoint and DNA repair^25^. Here, we not only demonstrate a novel function of CHK1 in regulation of TOP1cc dynamics (which may contribute to cancer cell proliferation to impart therapeutic resistance), we also unravel an additional layer of complexity in the molecular mechanism of action of CHK1 inhibitors. These newly uncovered aspects bear critical importance in further understanding of the complex mechanisms of regulation of TOP1 as well as the roles of played by CHK1 in ensuring integrity of the human genome. Our study also highlights that CHK1*i*-mediated sensitization of cancer to various therapeutics cannot not be attributed solely to defective cell cycle checkpoints and inhibition of DNA repair, but also to the accumulation of TOP1cc-mediated impaired replication and generation of replication-transcription conflicts, R-loop, DSBs and chromosomal instability.

In summary, our study uncovers three important aspects of CHK1 and TOP1 biology (**Figure 7E**). Firstly, it establishes a hitherto-unknown role of CHK1 in suppressing genomic instability through direct regulation of TOP1 catalytic activity, thus widening its ambit as a protector of genomic integrity. Secondly, it adds a layer of complexity to the established paradigm of the mechanism of action of CHK1 inhibitors. Thirdly, failure of the PTM-proteasome-nucleolytic machinery to identify CHK1*i*-mediated TOP1ccs for longer duration opens up a new attractive therapeutic window which bears the potential to evade the degradation of TOP1ccs in response to classical TOP1 poisons.

## Materials and Methods

### Chemicals and Antibodies

Unless otherwise stated, all fine chemicals were purchased from Sigma Aldrich (St Louis, MO, USA). SCH900776, UCN-07, NU7026, PHA-767491 and AZD6738 were purchased from Selleck Chemicals (Houston, TX, USA). All other kinase inhibitors were purchased from Cayman Chemical Company (Ann Arbor, MI, USA). Rabbit polyclonal anti-TOP1 antibody (#HPA019039), rabbit polyclonal anti-GFP antibody (#G1544), mouse monoclonal anti-DNA-RNA hybrid antibody (#MABE1095), mouse monoclonal anti-TOP1cc antibody (#MABE1084), mouse monoclonal anti-DNA antibody (#MAB030), mouse monoclonal anti-phospho-Histone H2AX (Ser139) clone JBW301 antibody (#05-636-I), rabbit monoclonal anti-53BP1 antibody (#ZRB1153), mouse monoclonal anti-ubiquitin antibody (#ST1200), mouse monoclonal anti poly-ADP-ribose antibody (#MABC547) were purchased from Sigma Aldrich (St Louis, MO, USA). Mouse monoclonal anti-CHK1 antibody (#sc-8408) was purchased from Santa Cruz Biotechnology (Dallas, TX, USA). Rabbit polyclonal anti-SUMO1 (#4930) and rabbit monoclonal anti-SUMO2/3 (#4971) antibodies were purchased from Cell Signalling Technology (Danvers, MA, USA). All AlexaFluor tagged secondary antibodies were purchased from Jackson Immunoresearch Laboratories (West Grove, PA, USA).

Antibodies used for DNA fiber assay were as follows: BrdU (IdU) antibody (#347580) from BD Pharmingen (CA, USA); BrdU (CldU) (#ab6326) from Abcam (MA, USA), Cy3-conjugated anti-Rat IgG (#712-165-153) and Alexa flour 488-conjugated anti-Mouse IgG (#715-546-151) from Jackson Immunoresearch Laboratories (West Grove, PA, USA). EdU and EU incorporation kits (#C10337, #C10339 and #C10330) were purchased from Thermo Fisher Scientific (Waltham, MA, USA). Duolink Proximity Ligation Assay components (#DUO92002, DUO92004 and DUO92008) were purchased from Sigma Aldrich (St Louis, MO, USA). SsoAdvanced Universal SYBR Green supermix was purchased from Bio-Rad Laboratories (Hercules, CA, USA). GFP trap magnetic agarose was purchased from Proteintech Group (Rosemont, IL, USA). Purified human TOP1 and CHK1 proteins were purchased from TopoGEN (Buena Vista, CO, USA) and Abcam (Cambridge, UK). ^32^P-ATP was purchased from Board of Radiation and Isotope Technology, India.

### Cell culture

U2-OS, PANC-1, A549 and HCT116 cells were procured from ECACC. MDA-MB-231 cells were procured from ATCC. Cells were cultured in Dulbecco’s Modified Eagle Medium supplemented with 10% fetal bovine serum and Penicillin-Streptomycin-Amphotericin B mix. Cells were maintained at 37 ℃ under 95 % relative humidity and 5% CO_2_.

### RADAR Assay

<u>R</u>apid <u>A</u>pproach to <u>D</u>NA <u>A</u>dduct <u>R</u>ecovery (RADAR) assay^50^ was performed to quantify cellular levels of TOP1ccs. Briefly, 1.5 × 10^5^ cells were seeded per well in 6 well plates (to achieve ∼90 % confluence), and allowed to attach for 16 hours. Cells were then subjected to relevant treatments. Post treatment, cells were washed once with phosphate buffered saline (PBS), followed by lysis with DNAzol reagent. DNA was then precipitated with one volume of absolute ethanol, recovered by spooling, and washed once with 80% ethanol. Pellets were briefly air dried for 2-3 minutes, followed by resuspension in 8 mM NaOH. DNA was diluted in Tris Buffered Saline, followed by quantification. DNA (500 ng per sample) was then spotted on nitrocellulose membrane using a slot blot apparatus (BioRad, USA) under vacuum. The membrane was air dried, sandwiched in Whatman Paper and baked at 80 ℃ for 2 h in a hot air oven. Post baking, membrane was blocked for 1 h in 2% BSA in PBS-Tween 20. Membrane was then incubated overnight at 4 ℃ with primary antibodies against TOP1 and DNA, respectively, followed by 2 h incubation with requisite secondary antibodies at room temperature. Blots were developed using WesternBright Sirius HRP ECL substrate (Advansta, CA, USA).

### RADAR-based screening of kinase inhibitors

A manually curated library of 25 small molecule kinase inhibitors was employed to evaluate their ability to induce TOP1ccs. Briefly, 7.5 × 10^4^ cells were seeded per well in 12 well plates. Cells were treated with relevant concentrations of inhibitors for 2, 4 or 6 h, followed by sample processing as per RADAR assay protocol. Densitometric analysis of blots was used for comparative evaluation of TOP1cc stabilization. All signals were normalized to the corresponding DNA control. Relative TOP1cc stabilization under each treatment condition was calculated as fold change over untreated condition.

### Fluorescence Recovery After Photobleaching (FRAP)

FRAP-mediated quantification of TOP1 dynamics was performed as described previously^7^. Briefly, U2-OS cells stably expressing EGFP-TOP1 (1.5 × 10^5^ cells per dish) were seeded in thin-bottom confocal grade 35 mm dishes. After 16 h, cells were subjected to requisite treatments and observed under a Zeiss LSM 780 confocal microscope employing a 488 nm laser line. Image acquisition was performed at 40X optical and 6X digital magnification, ensuring a pixel dwell time of 1 µs. Five pre-bleach images were acquired at 1% laser power, followed by bleaching of the region of interest (ROI) with 20 iterations at 100% power. Post bleaching, 250 images were captured in a time series with 1 ms interval. A reference ROI was also included throughout the experiment to apply photobleaching-correction. Image analysis was performed using Zen 2.3 SP1 software to quantitatively determine the percentage immobile fraction. Additionally, percentage fluorescence recovery was determined according to the following formula-

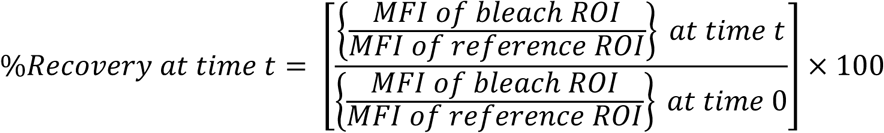

MFI represents Mean Fluorescence Intensity.

### Immunoblotting

Western blotting was performed as reported previously ^51^. Briefly, 0.8 × 10^6^ cells were seeded in 60 mm dishes and allowed to attach for 16 hours. Cells were subjected to treatment, washed once with PBS, and scraped using a policeman. The cell pellet was washed once with PBS, and lysed with an appropriate volume of RIPA buffer (10 mM Tris-HCl pH 8.0, 1 mM EDTA, 0.5 mM EDTA, 1% Triton X-100, 0.1% Sodium deoxycholate, 0.1% SDS, 140 mM NaCl) supplemented with protease inhibitor cocktail and 10 Units/ml DNase I. Cells were lysed on ice for 1 h, followed by centrifugation of lysate at 12000 rpm for 20 minutes. Total protein was quantified using Bradford assay. 50 µg protein was resolved on SDS-PAGE, followed by electrotransfer on methanol-activated PVDF membrane using a BioRad semi-dry transfer apparatus. Membrane was blocked for 1 h with 2 % BSA-PBST, followed by incubation with requisite primary and secondary antibodies.

### Assay for TOP1 downregulation

CPT or SCH induced TOP1 degradation was assayed using alkaline cell lysates as reported previously ^52^. Briefly, 1.5 × 10^5^ cells were seeded in 6 well plates, and subjected to relevant treatments 16 h post seeding. Post treatment, cells were washed once with PBS, followed by alkaline lysis (200 mM NaOH and 2 mM EDTA). Lysates were then collected and neutralized using 1 volume of Tris pH 8.0 (1 M). Appropriate volumes of 10X DNase I digestion buffer and 50X protease inhibitor cocktail (Roche) were added to achieve 1X final concentration of each. DNA was digested on ice for 1 h using a cocktail of DNase I and Benzonase. Samples were then boiled in SDS sample buffer for 20 minutes and subjected to western blotting for TOP1.

### DUST Assay

<u>D</u>etection of <u>U</u>biquitylated and <u>S</u>UMOylated <u>T</u>OP1ccs (DUST) assay was performed, as described elsewhere^13^. Briefly, 1.6 × 10^6^ cells were seeded in 90 mm dishes (to achieve ∼90% confluence) and allowed to attach for 16 h. Cells were subjected to relevant treatments, washed once with PBS, and lysed with 1 mL DNAzol reagent. DNA was precipitated with one volume of absolute ethanol, recovered by spooling and washed once with 70% ethanol. Pellets were briefly air dried, resuspended in Tris-EDTA (TE) buffer (pH 8.0), and heated at 65 ℃ for 15 minutes. DNA was then sheared on ice using a Cole-Parmer Ultrasonic Processor (40 % amplitude; 4 cycles; 10 seconds on/10 seconds off). Samples were then centrifuged at 12500 rpm for 5 minutes, followed by RNase A treatment of supernatant for 1 h at 4 ℃. DNA was then precipitated overnight at -20 ℃, using 1/10^th^ volume of 3 M sodium acetate and 2.5 volumes of absolute ethanol. DNA was recovered by centrifugation, washed once with 80 % ethanol, and resuspended in 100 µl of TE buffer. DNA was then quantified, and 25 µg DNA per sample was digested for 2 h at 37 ℃ using a mix of 10 units of DNase I and 10 units Benzonase. Digested samples were boiled with SDS sample buffer for 10 minutes, followed by SDS-PAGE on 6 % polyacrylamide gels. SUMOylation, PARylation, and Ubiquitination was detected using specific antibodies. Additionally, 2 µg DNA per sample was subjected to dot blotting and subsequent immunodetection using anti-dsDNA antibody to serve as a loading control.

### EdU and EU incorporation assays

Click-EdU and Click-EU assays were employed for determining levels of cellular replication and transcription, respectively. Cells were seeded on coverslips, followed by appropriate treatments. 30 minutes prior to termination of experiment, cells were incubated with 10 µM EdU or 1 mM EU. Cells were then washed with PBS and fixed with 4% paraformaldehyde for 15 minutes at 4 ℃. Fixed cells were permeabilized with PBS supplemented with 0.25% v/v Triton X-100, and Click reaction was subsequently performed using Click-IT kit (Invitrogen, USA) according to the manufacturer’s instructions. Alexa Fluor 488 or Alexa Fluor 594 azide was used as per the experimental requirements. Post Click reaction, coverslips were washed once with PBS, air dried, and mounted mounted using 80% glycerol supplemented with 10 µg/mL Hoechst 33258. Cells were then observed under a Zeiss LSM 780 confocal microscope.

### Immunofluorescence

Immunofluorescence assays were performed as described previously^52^. Briefly, cells were seeded on coverslips, followed by requisite treatments 16 h post seeding. Cells were washed once with PBS and fixed with 4% paraformaldehyde for 15 minutes at 4 ℃. Cells were permeabilized with PBS supplemented with 0.25 % v/v Triton X-100. Cells were washed once with PBS, and blocked for 1 h at room temperature in 1% BSA in PBS. Coverslips were incubated with relevant primary antibodies (diluted in 2.5 % BSA in PBS) overnight at 4 ℃, followed by 3 washes (5 minutes each) with PBS. Cells were then incubated with Alexa Fluor 488 or Alexa Fluor 594 tagged secondary antibodies at room temperature for 2 hours under subdued light and washed thrice with PBS (5 minutes each). Coverslips were then air-dried and mounted on glass slides using 80% glycerol supplemented with 10 µg/ml Hoechst 33258. In case of experiments involving simultaneous immunofluorescence and EdU incorporation, Click EdU reaction was performed at the end of secondary antibody incubation.

### TOP1cc Immunofluorescence

Immunofluorescence-based detection of TOP1ccs was performed as described previously^52^. Briefly, 1.5 × 10^5^ cells were seeded on coverslips in 6 well plates. Cells were subjected to treatment 16 hours post seeding. Coverslips were washed and fixed with 4% paraformaldehyde for 15 minutes at 4 ℃. Cells were then permeabilized with PBS-0.25 % Triton X-100 for 15 minutes at 4 ℃. Permeabilization was followed by denaturation with 2% SDS for 10 min at 20 ℃. Cells were washed thrice with PBS (5 min each), followed by blocking with PBS supplemented with 0.05 % Tween 20, 0.01% Triton X-100 and 1% BSA (PBSTT) at room temperature for 2 h. Coverslips were then incubated with primary antibody overnight at 4℃. Coverslips were then washed thrice with PBSTT (5 min each), and incubated for 2 h at room temperature with secondary antibody under subdued light. Post incubation, coverslips were washed thrice with PBSTT (5 min each), air dried, and mounted with 80% glycerol supplemented with 10 µg/mL Hoechst 33258.

### Cell counting experiments for comparative evaluation of cell growth kinetics

In order to compare the growth kinetics of U2-OS^EGFP-TOP1WT^ and U2-OS^EGFP-TOP1S320A^ stable cells, 4000 cells were seeded in 60 mm dishes. Cells were recovered through trypsinization 3, 6 and 9 days post seeding, and cell number was enumerated using a hemocytometer.

### RNA/DNA hybrid immunofluorescence

Microscopic detection of RNA-DNA hybrids was performed as described elsewhere^53^ with minor modifications. Briefly, exponentially growing cells were seeded on coverslips and allowed to attach for 16 hours. Cells were fixed with 4% paraformaldehyde (in PBS) at room temperature for 10 min. Cells were then permeabilized with 0.5 % ice-cold Triton X-100. Cells were then subjected to RNase A pre-treatment in RNase A digestion cocktail (6 µg/mL RNase A in 10 mM Tris-HCl pH 7.5 and 500 mM NaCl) for 45 minutes at 37 ℃. For RNase H control, cells were subjected to co-digestion with RNase A and RNase H. Post digestion, cells were blocked with 5 % BSA in PBS for 3 h at room temperature. Cells were then incubated with anti-DNA-RNA hybrid (S9.6) antibody (diluted in 2.5 % BSA-PBS) overnight at 4 ℃. Cells were then washed thrice with PBS, followed by incubation with secondary antibody (diluted in 2.5 % BSA-PBS) for 2 h under subdued lighting. Coverslips were washed thrice with PBS, air-dried, and mounted on slides with 80% glycerol supplemented with 10 µg/mL Hoechst 33258.

### Detection of RNA/DNA hybrids through dot blot

For dot blot-mediated detection of RNA-DNA hybrids, cells were lysed with DNAzol reagent, followed by precipitation on genomic DNA with 1 volume of 100% ethanol. Precipitated genomic DNA was washed once with 70% ethanol, followed by resuspension in 1X NEB CutSmart buffer. Genomic DNA was randomly fragmented overnight with a cocktail of restriction enzymes consisting of EcoRI, HindIII, XbaI and NotI. Fragmented DNA was subjected to extraction with phenol-chloroform-isoamyl alcohol (25:24:1). DNA was then precipitated overnight at -20 ℃ with 3M sodium acetate (1/10^th^ volume) and 100% ethanol (2.5 volumes). DNA pellet was washed once with 70% ethanol, and resuspended in TE buffer. DNA was then quantified, and 1 µg of DNA was loaded per sample. Further processing was similar to RADAR assay. For determining RNase H sensitivity of signal, 1 µg of DNA sample was digested with RNase H for 6 h prior to loading.

### Plasmids and transfections

EGFP-TOP1^WT^ plasmid was a kind gift from Professor Benu Brata Das, Indian Association for the Cultivation of Science, Kolkata, India. Transfections were carried out using Lipofectamine 2000 reagent (Invitrogen, USA) according to the manufacturer’s protocol. A DNA:Lipofectamine ratio of 1:1.5 was maintained for all transfections. For immunoprecipitation experiments, transfections were performed using the calcium phosphate method.

### Site-directed mutagenesis

Site directed mutagenesis was performed as described previously^7^. The following primers were used: S320A forward 5’-GAAACAGATGGCCAAGGAAGAGA-3’; reverse 5’-GAAACAGATGGCCAAGGAAGAGA-3’. Mutations were confirmed through sequencing.

### Mass Spectrometric analysis of TOP1 phosphorylation sites

A mass tandem mass spectrometry based assay was employed to detect amino acids phosphorylated on catalytically active TOP1. Briefly, exponentially growing cells were seeded in 90 mm dishes to achieve ∼90% confluence ∼16 h post seeding. To specifically enrich catalytically active TOP1, cells were treated for 30 mins with 1 µM CPT to trap TOP1 molecules actively involved in catalysis on DNA. Cells were then disrupted with DNAzol reagent, followed by downstream processing of samples as in case of DUST assay. An excess of untreated sample was also included (to recover enough TOP1 protein). Protease and phosphatase inhibitors were included in all steps post lysis. Samples were resolved on SDS PAGE, followed by gel excision in the 100-125 KDa region. Further sample processing and analysis was performed (Valerian Chem Private Limited, India). Briefly, Gel bands were cut into small pieces and reduced with 5 mM tris(2-carboxyethyl)phosphine (TCEP) followed by alkylation with 50 mM iodoacetamide. Samples were then digested with Trypsin (1:50, Trypsin/lysate ratio) for 16 h at 37 °C. Digests were cleaned using a C18 silica cartridge to remove salts and vacuum dried. Dried pellets were resuspended in buffer A (5% acetonitrile, 0.1% formic acid). All analyses were performed on an Ultimate 3000 RSLCnano system coupled with an Orbitrap Eclipse. Approximately 500 ng of each sample was loaded on C18 column (50 cm, 3.0 μm Easy-spray column, Thermo Fisher Scientific). Peptides were eluted with a 0–40% gradient of buffer B (80% acetonitrile, 0.1% formic acid) at a flow rate of 300 nl/min and injected for MS analysis. LC gradients were run for 100 minutes. MS1 spectra were acquired in the Orbitrap (R= 240k; AGQ target = 400 000; Max IT = 50 ms; RF Lens = 30%; mass range = 400−2000; centroid data). Dynamic exclusion was employed for 10 s excluding all charge states for a given precursor. MS2 spectra were collected in linear ion trap (rate = turbo; AGQ target = 20 000; MaxIT = 50 ms; NCE_HCD_= 35%).

RAW files generated were analyzed with Proteome Discoverer (v2.2) against the single reference proteome database. For Sequest search, the precursor and fragment mass tolerances were set at 10 ppm and 0.8 Da, respectively. The protease used to generate peptides, *i.e.* enzyme specificity was set for trypsin/P (cleavage at the C terminus of “K/R: unless followed by “P”) along with maximum missed cleavages value of two. Carbamidomethyl on cysteine as fixed modification and oxidation of methionine and N-terminal acetylation and Phospho (S,T,Y) were considered as variable modifications for database search. Both peptide spectrum match and protein false discovery rate were set to 0.01 FDR.

### Isolation of mouse splenocytes

Splenic cells were isolated from BALB/c mice. Briefly, 4-6 weeks old male mice were euthanized and spleen was harvested under aseptic conditions. The spleen was sliced into small pieces and thoroughly washed thrice with ice cold PBS. A single cell suspension was created by passing the excised spleen pieces through a cell strainer with the help of the plunger of a sterile 1 mL syringe. The cell suspension was centrifuged at 1600 rpm for 5 min, and the pellet was resuspended in 2 mL of pre warmed RBC lysis buffer (193 mM NH_4_Cl, 12.5 mM KHCO_3_, 0.01 % w/v EDTA). The cell suspension was incubated at 37 ℃ for 2 minutes. Post incubation, 30 mL of PBS was added and the cells were centrifuged at 1600 rpm for 5 minutes. The cell pellet was resuspended in appropriate volume of PBS.

### RNA Sequencing and Data Analysis

RNA sequencing was performed through contract research. Briefly, cells were seeded in 60 mm dishes, and subjected to requisite treatments 16 h post seeding. Cells were then washed twice with ice cold PBS, and harvested through trypsinization. Total RNA was isolated using Qiagen RNeasy Mini Kit according to the manufacturer’s instructions. Paired end sequencing (average read length ∼150 base pairs) was performed through Illumina Sequencing by Synthesis, and a total of ∼50 million reads were analysed per sample. Differential gene expression analysis (with respect to untreated sample) was performed through contract research, and downstream data analysis was performed as follows. Downregulated genes were selected with a fold change cutoff of 0.5. Unaltered genes were defined as genes with a fold change lying between 0.9 and 1.1. Gene lengths were retrieved from NCBI through deployment of a python code (reproduced in supplementary methods) run on Google Colaboratory environment.

### *In vitro* kinase assay

*In vitro* kinase assay was performed with purified, catalytically active human CHK1 and TOP1 proteins in kinase reaction buffer (HEPES pH 7.5, 1 mM DTT, 100 µM Na_3_VO_4_, 10 mM MgCl_2_). Reactions were initiated by adding 50 µM ATP (spiked with 10 µCi ^32^P-labelled ATP) to CHK1 and TOP1 (2 picomoles each). Samples were incubated at 30 ℃ for 30 minutes. Reactions were terminated by adding SDS sample buffer. Samples were boiled at 95 ℃ for 10 minutes, followed by resolution on SDS PAGE. Gel was dried, and subjected to autoradiography.

### *In vitro* plasmid relaxation assay

*In vitro* plasmid relaxation assay^52^ was performed in TOP1 reaction buffer (10 mM Tris-Cl pH 7.9, 1 mM EDTA, 150 mM NaCl, 0.1% BSA, 0.1 mM Spermidine, 5% glycerol). Supercoiled pUC19 plasmid DNA was incubated with TOP1 for varying lengths of time at 37 ℃, and reactions were terminated using 5X Stopping Buffer (5% sarkosyl, 0.0025% bromophenol blue, 25% glycerol). Samples were resolved on a 1 % agarose gel in TAE buffer at 2.5 V/cm for 6 h. Gel was washed once with deionized water, followed by staining with 1 µg/mL ethidium bromide.

### Preparation of crude nuclear extracts

Total cellular TOP1 activity was assayed as reported elsewhere^54^ with minor modifications. Briefly, exponentially growing cells (5 × 10^6^ cells per 150 mm dish) were subjected to relevant treatments, washed once with ice cold PBS, and harvested by scraping. The pellet was resuspended in 180 µL of ice cold low salt extraction buffer (20 mM Tris-HCl pH 7.5, 5 mM KCl, 1 mM MgCl_2_, 10 % v/v glycerol, protease inhibitor cocktail) and incubated on ice for 10 minutes. Cells were subjected to 15-20 strokes of a pre-cooled Dounce homogeniser, followed by 30 min; incubation on ice. Nuclei were recovered by centrifugation at 12500 rpm for 3 minutes at 4 ℃. The cytosolic extract was discarded, and the nuclear pellet was resuspended in 180 µL of ice cold high salt extraction buffer (low salt extraction buffer with 350 mM KCl, supplemented with protease inhibitor cocktail), followed by 80 min incubation on ice. The nuclear extract was clarified by centrifugation at 12500 rpm for 10 minutes at 4 ℃. Total protein was quantified using Bradford method, followed by *in vitro* plasmid relaxation assay with different amounts of nuclear extract. For SCH treated samples, low salt and high salt extraction buffers were supplemented with SCH (25 nM) to prevent CHK1 reactivation during lysis.

### Co-immunoprecipitation

For co-immunoprecipitaion experiments, 5 × 10^6^ cells were seeded per dish in 150 mm dishes. Cells were transfected with 15 µg of EGFP-TOP1 plasmid 16 hours post transfection by calcium chloride method. The cell medium was replaced with fresh medium 24 h post transfection. Cells were incubated with the fresh medium for 24 h, followed by washing with ice cold PBS, and harvesting by scraping. Cell pellets were washed once with PBS, and lysed with IP lysis buffer (50 mM Tris-HCl pH 8.0, 300 mM NaCl, 0.4 % NP-40, protease inhibitor cocktail), supplemented with DNase I digestion buffer (1X final concentration from a 10X stock), 10 U/mL DNase I and 5 U/mL benzonase. Cells were lysed on ice for 1 h with periodic voxtexing. Cell lysates were centrifuged at 12500 rpm for 20 minutes to eliminate cellular debris. The supernatants were collected and diluted to 1 mL volume with IP dilution buffer (50 mM Tris-HCl pH 8.0, 0.4 % NP-40, protease inhibitor cocktail). Total protein was quantified by Bradford method, and 5 % of each lysate was saved as input. Further, 1 mg protein per sample was incubated with anti-GFP antibody overnight at 4 ℃ on a rotary shaker. Protein A/G magnetic agarose beads were equilibrated with IP dilution buffer and 25 µl beads were added per sample and incubated for 4 hours at 4 ℃ on a rotary shaker. Beads were then washed, boiled with SDS sample buffer and samples were analysed by western blotting.

### Immunoprecipitation of active wild type and mutant TOP1 from human cells

For immunoprecipitation of TOP1, 5 × 10^6^ cells were seeded in 150 mm dishes. Cells were allowed to attach for 16 hours and transfected with 15 µg each of EGFP-TOP^WT^ or EGFP-TOP1^S320A^ plasmid by calcium chloride method. Cell culture medium was replaced 24 h post transfection; 24 h later, cells were washed once with ice cold PBS, and harvested with a policeman. Cell pellets were washed once with PBS, followed by lysis in IP lysis buffer (10 mM Tris-HCl pH 7.5, 150 mM NaCl, 0.5 mM EDTA, 0.5 % NP-40). IP lysis buffer was supplemented with protease inhibitor cocktail, DNase I digestion buffer (1X from a 10X) stock, 10 U/mL DNase I and 5 U/mL benzonase. Lysis was performed on ice for 1 h with intermittent vortexing. Cellular debris was precipitated by centrifugation at 12500 rpm for 20 min. The cell lysates were diluted to 1 mL volume in IP dilution buffer (10 mM Tris-HCl pH 7.5, 150 mM NaCl, 0.5 mM EDTA) supplemented with protease inhibitor cocktail. Lysates were subjected to total protein quantification (Bradford method), and 1 mg lysate per sample was subjected to immunoprecipitation overnight using GFP-Trap Magnetic Agarose (ChromoTek GmBh) at 4 ℃ on a rotary shaker. 5 % volume of each sample was reserved as input. Beads were then washed thrice with IP wash buffer (10 mM Tris-HCl pH 7.5, 150 mM NaCl, 0.5 mM EDTA, 0.05 % NP-40) supplemented with protease inhibitor cocktail. Beads were resuspended in 50 µl of TOP1 reaction buffer (100 mM Tris-Cl (pH 7.9), 10 mM EDTA, 1.5 M NaCl, 1% BSA, 1 mM Spermidine, 50% glycerol). For verification of immunoprecipitation efficiency, 10 µl of beads were removed and boiled with SDS sample buffer, followed by western blotting with anti-GFP antibody. The remaining beads were used for plasmid relaxation assays.

### Chromatin immunoprecipitation

For chromatin immunoprecipitation, 5 × 10^6^ cells were seeded in 150 mm dishes. Cells were transfected with requisite plasmids by calcium chloride method (described in plasmids and transfections section). 48 h post transfection, cells were harvested by scraping, followed by one wash with ice cold PBS. Cell pellet was then resuspended in ice-cold swelling buffer (25 mM HEPES pH 7.8, 1.5 mM MgCl_2_, 10 mM KCl, 0.1 % NP-40 and protease inhibitor cocktail), and incubated on ice for 10 minutes. Cells were then disrupted with 20 strokes of a Dounce homogenizer. Nuclei were precipitated at 2000 rpm for 5 min followed by resuspension in sonication buffer (50 mM HEPES pH 7.8, 140 mM NaCl, 1 mM EDTA, 1% Triton X-100, 0.1 % sodium deoxycholate, 0.1% SDS, protease inhibitor cocktail), sonicated on ice for 18 cycles (10 s on/30 s off; 50% amplitude) with a Cole-Parmer ultrasonic processor. Samples were then centrifuged at 13000 rpm for 15 minutes at 4 ℃. The supernatant was recovered, and total DNA was spectrophotometrically quantified. For immunoprecipitation, 100 µg of genomic DNA per sample was incubated overnight with pre-equilibriated GFP-Trap Magnetic Agarose beads at 4 ℃. Further, 5% DNA from each samples was reserved for preparation of input sample. Beads were then subjected to two washes each with sonication buffer, wash buffer A (50 mM HEPES pH 7.8, 500 mM NaCl, 1 mM EDTA, 1 % Triton X-100, 0.1 % sodium deoxycholate, 0.1% SDS, protease inhibitor cocktail), and TE buffer (10 mM Tris-HCl pH 8, 1mM EDTA), respectively. Beads were then treated with 200 µL elution buffer (50 mM Tris pH 8, 1 mM EDTA, 1% SDS, 50 mM sodium bicarbonate) for 15 min at 65 ℃. Elution was performed twice, and the eluates were combined. The volumes of input samples were made up to 400 µL with elution buffer. 21 µL NaCl was added to each sample, and 1 µL RNase A was added (from a 10 mg/mL stock). Samples were digested for 1 h at 37 ℃. Samples were supplemented with 20 µL EDTA (from a 100 mM stock) and digested with 2 µL Proteinase K (from a 10 mg/mL stock) for 2 h at 42 ℃. Samples were extracted once with phenol-chloroform-isoamyl alcohol (25:24:1) and once with chloroform-isoamyl alcohol (24:1), followed by precipitation overnight with 1/10^th^ volume sodium acetate and 2.5 volumes of 100 % ethanol. Glycogen (1 µL from a 20 mg/mL stock) was included as a carrier. Samples were centrifuged at 13000 rpm for 30 min, washed once with 70% ethanol, and resuspended in an appropriate volume of 10 mM Tris-HCl pH 8.

Quantitative real-time PCR was performed using SYBR green method. Details of primers employed are provided in Supplementary Table 1. All quantifications were made in terms of percentage of input, and relative ChIP signal was determined based on β-Actin TSS of EGFP-TOP1^WT^ sample, which was set to 1.

### DNA/RNA hybrid immunoprecipitation (DRIP)

For immunoprecipitation of RNA/DNA hybrids, nuclei were isolated as described for immunoprecipitation. Isolated nuclei were lysed on ice with nuclear lysis buffer (50 mM Tris-HCl pH 8.0, 5 mM EDTA, 1% SDS) for 30 minutes. Nuclear lysates were digested with Proteinase K for 3 h at 55 ℃. Genomic DNA was then precipitated with 100 % ethanol, washed once with 70 % ethanol and resuspended in IP dilution buffer (16.7 mM Tris-HCl pH 8.0, 1.2 mM EDTA, 167 mM NaCl, 0.01% SDS, 1.1% Triton X-100). Genomic DNA was sonicated as described for chromatin immunoprecipitation. Sonicated DNA was quantified, and 100 µg DNA was incubated overnight with 4 µg of S9.6 antibody and a 1:1 mix of pre-equilibriated Protein A/Protein G magnetic beads at 4℃. In addition, 5 % DNA was stored as input. Post incubation steps were identical to those described for chromatin immunoprecipitation. To test the RNase H sensitivity of the signal, samples were treated with 2 U of RNase H before being subjected to immunoprecipitation. Appropriate isotype control was also included. Determination of relative DRIP signal was identical to chromatin immunoprecipitation.

### Metaphase Preparations

Metaphase spreads were prepared as described elsewhere ^55^ with minor modifications. Briefly, cells were grown in 60 mm dishes to sub-confluence. Cells were then incubated in fresh culture medium supplemented with 200 ng/mL nocodazole for 16 hours. Cells were then harvested through mitotic shake-off and washed once with pre-warmed PBS. Cells were then resuspended in swelling buffer (75 mM KCl) and incubated at 37 ℃ for 6 minutes. Cells were centrifuged at 1800 rpm, for 10 minutes, followed by resuspension in fixative solution containing methanol and acetic acid in a ratio of 3:1. The fixation step was repeated twice, followed by resuspension in 200 µL of fixative. Cells were dropped on chilled, pre-hydrated glass slides, and dried immediately on a hot plate. The dried slides were immersed in a coplin jar containing 5 % solution of Giemsa stain in phosphate buffer. Slides were stained for 20 min, followed by thorough washing with nanopure water. Images were captured with a light microscope at 100x magnification. A minimum of 100 metaphase spreads were analysed per sample.

### DNA fiber analysis

DNA fiber analysis was performed as described elsewhere ^56^ with minor modifications. Briefly, 2×10^5^ cells were plated per well in 6 well plates and transfected with EGFP-TOP1^WT^ and EGFP-TOP1^S320A^ with Lipofectamine 3000 reagent according to manufacturer’s instructions. 48 h post transfection, cells were sequentially labelled with 30 µM CldU and 300 µM IdU (20 mins each) with 3 washes with excess volume of pre-warmed PBS in between. Cells were harvested post labelling, counted, and resuspended at a concentration of 1×10^6^ cells/mL. Labelled and unlabelled cells were mixed at a ratio of 1:4, and 2.5 µL of the cell suspension was deposited on a glass slide. This was followed by addition of 6 µL of lysis buffer (0.5 % SDS, 200 mM Tris-HCl pH704 and 50 mM EDTA). The slide was incubated for 6 min at room temperature followed by tilting the slide at ∼30°, allowing the droplet to spread down the slide under gravity. The slide was dried, followed by fixation for 15 min with methanol-acetic acid mixture (3:1). Slides were incubated in 100% methanol for 5 min, followed by two washes with PBS. DNA was denatured for 1 h in 2.5 N HCl. Slides were washed thrice with PBS, followed by blocking (with 5 % BSA in PBS) for 45 mins at 37℃. Slides were washed once with PBS, followed by treatment with primary antibodies (rat anti-CldU/BrdU and mouse anti-IdU/BrdU respectively) for 1 h at room temperature. Slides were washed thrice with PBS, followed by incubation with secondary antibodies for 1 h at room temperature. Slides were washed thrice with PBS and mounted in 80 % glycerol. Samples were visualised with a Leica MICA system under 63x water immersion objective. Image analysis was performed with ImageJ.

### Alphafold Multimer (AF-M) predictions

Predictions for CHK1-TOP1 interaction were made employing Alphafold Multimer. Data analysis was performed on the Cosmic^2^ server (https://cosmic2.sdsc.edu:8443/gateway/). Predictions were performed using Colabfold, employing alphafold2_multimer_v3 ^57,58^. The employed settings were as follows: number of models 5, number of recycles 3, Stop at score (Compute recycle models until plddt (single chain) or ptm score (complex) > threshold is reached.) 80. pdb100 was used as a template of published PDB structures. For TOP1, amino acids 201-765 were used for predictions. For CHK1, the following regions were employed: full length (1-476), Kinase domain (1-265) and KA1 domain (391-476). Model visualizations and analyses of pLDDT, pTM, iPTM and PAE values were performed using PAE Viewer webserver (http://www.subtiwiki.uni-goettingen.de/v4/paeViewerDemo)^59^. Analyses of pDOCKQ and SPOC scores were performed on the Predictome webserver (https://predictomes.org/)^60^.

### Statistical analysis

For statistical analysis, three or more independent experiments were performed in triplicates. Statistical analyses of above data were performed using Graphpad Prism 8. For DNA fiber analysis, Kruskal–Wallis test with Dunnett’s multiple comparisons was performed.

## Acknowledgement

The work has been funded by the Department of Atomic Energy, Government of India. The authors thank Dr. Benubrata Das for kindly providing EGFP-TOP1 expression construct. The authors also acknowledge the technical expertise provided by Dr. Usha Yadav for preparing metaphase spreads.

## Author contributions

AGM and BSP conceptualized, designed and executed the investigations. AGM has carried out most of the experiments. NC and PG carried out and validated few experiments. AGM, MS and BSP analysed and interpreted the data. AGM drafted the manuscript. MS and BSP revised the manuscript. All authors reviewed and approved the final version of the manuscript.

