## Supplementary Information for "CHK1-mediated regulation of TOP1 catalytic activity suppresses replication and transcription-associated genomic instability"

FIGURE S1

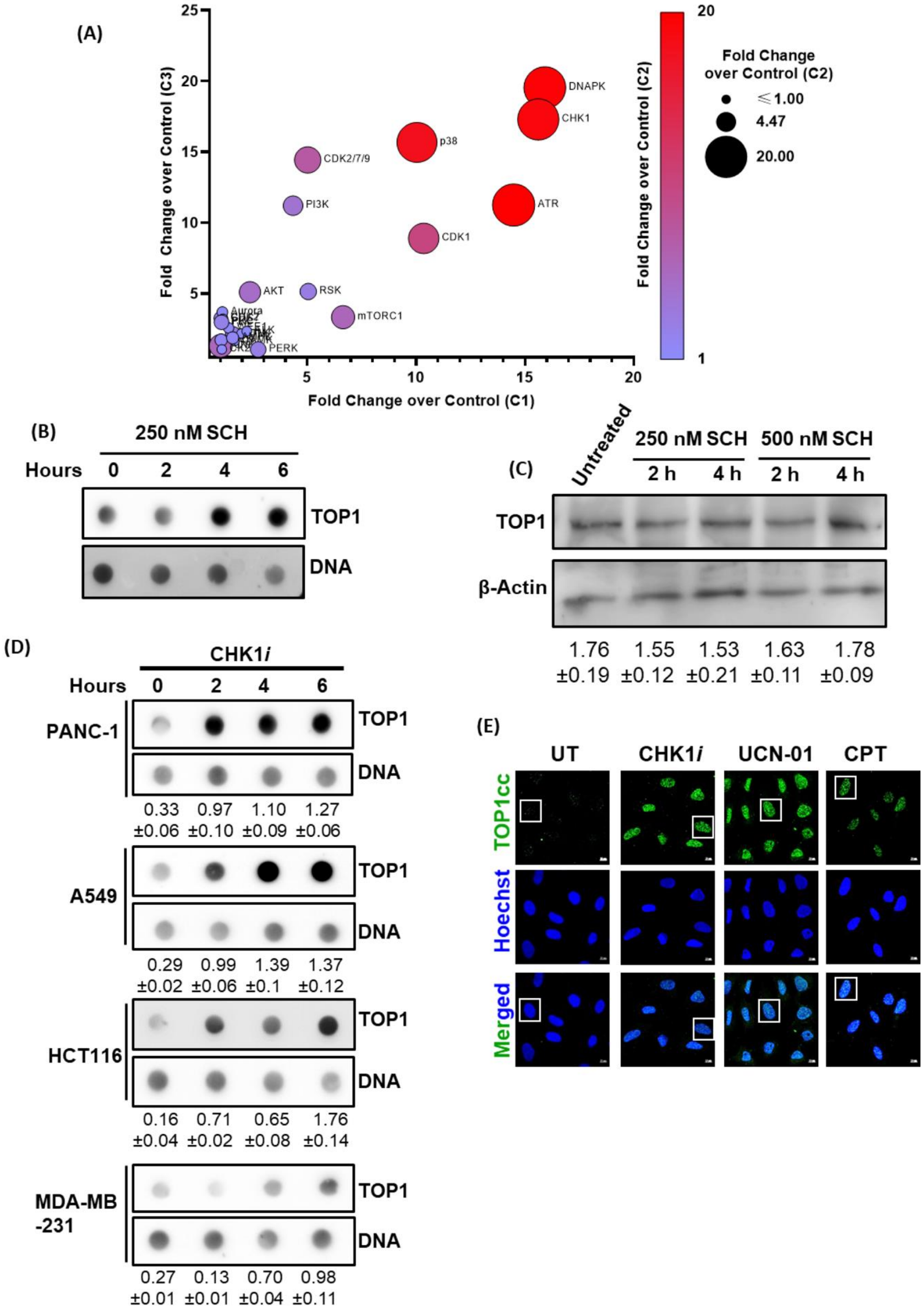


**Figure S1.** (A) A comprehensive representation of data derived from the kinase screen. Based on densitometric analysis of RADAR assay results, fold change relative to the control at the lowest (C1) and highest (C3) concentrations of each inhibitor was plotted along the X and Y-axes, respectively. The size as well the colour of the dots was used to represent the fold change at the intermediate (C2) concentration. At each concentration, the maximum fold change value was considered irrespective of the timepoint. (B) Time dependent TOP1cc stabilization in U2-OS cells in response to CHK1*i* treatment. U2-OS cells were treated with 250 nM CHK1*i* for 2, 4 or 6 h, followed by RADAR assay. (C) Estimation of global TOP1 levels in U2-OS cells treated with SCH. U2-OS cells were treated with 250 or 500 nM SCH for indicated lengths of time followed by estimation of global TOP1 levels by the alkaline lysis method. (D) SCH-mediated TOP1cc stabilization in different cancer cells. MDA-MB-231, PANC-1, A549 and HCT116 cells were treated with 250 nM SCH for 2, 4 or 6 h, followed by RADAR assay. (E) TOP1cc immunofluorescence in U2-OS cells. U2-OS cells were treated with 250 nM SCH, 50 nM UCN-01 (2 h) or 1 µM CPT (1 h) followed by immunofluorescent detection with monoclonal anti-TOP1cc antibody. Scale bars: 10 µm.. Zoomed view image of nuclei, of same experiment, was presented in Figure 1Z2.

FIGURE S2

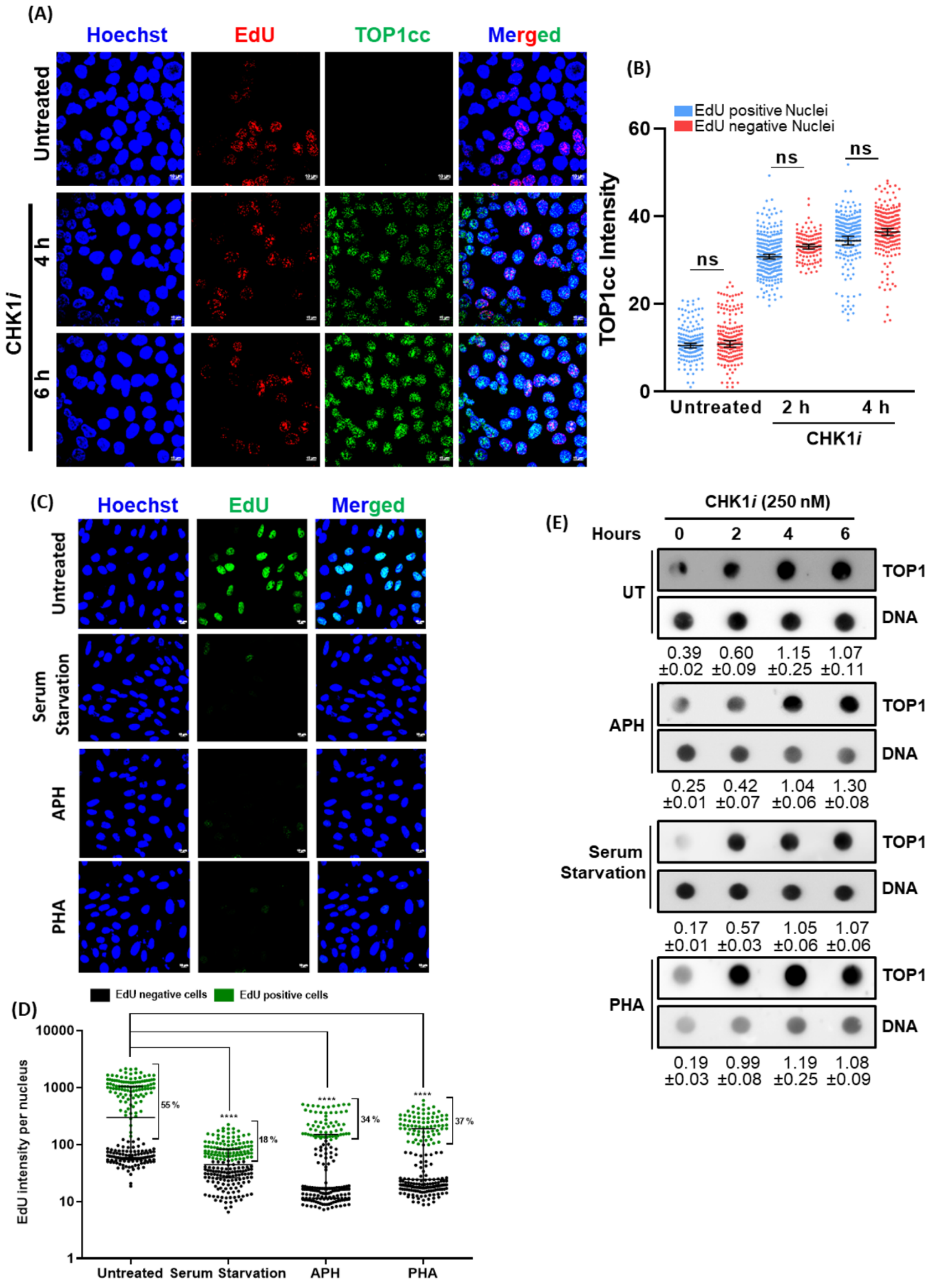


**Figure S2. CHK1*i* stabilizes TOP1ccs independent of replication and transcription.** (A, B) TOP1cc-EdU staining in U2-OS cells. U2-OS cells were treated with 250 nM CHK1*i* for 4

or 6 h, followed by 10 µM EdU treatment 30 min prior to termination of experiment. TOP1ccs were detected using anti-TOP1cc monoclonal antibody. (C, D) EdU incorporation in U2-OS cells under untreated, aphidicolin (APH; 250 nM, 16 h), serum starvation (0.5 % serum for 48 h) or CDC7i (PHA; 5 µM, 4 h) treatment conditions. U2-OS cells were subjected to requisite treatments, followed by incubation with 10 µM EdU 30 min prior to termination of experiment. EdU incorporation was measured using Click chemistry. (E) RADAR assay in U2-OS cells treated with CHK1*i* with or without pre-treatment with Aphidicolin (APH; 250 nM, 16 h pre-treatment), Serum starvation (0.5 % serum for 48 h), Cdc7i (PHA; 5 µM, 4 h pre-treatment). Scale bars: 10 µm. *ns* non-significant; *** $p$<0.001.

**FIGURE S3**

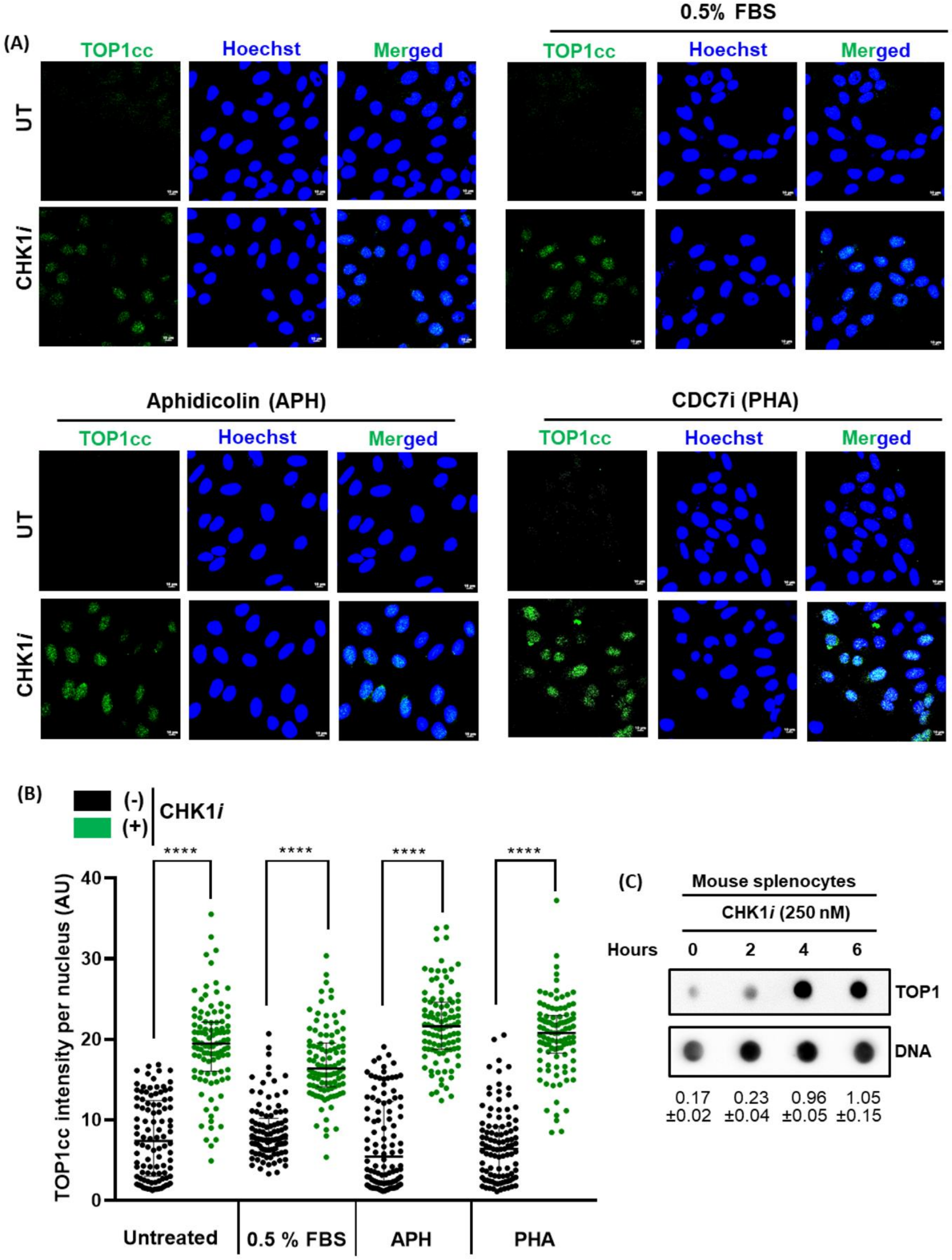


**Figure S3.** (A, B) Immunofluorescence mediated detection of TOP1cc levels in U2-OS cells subjected to CHK1*i*-treatment in the absence or presence of replication inhibition. U2-OS cells were subjected to serum starvation (0.5 % fetal bovine serum/FBS, 48 h), Aphidicolin (APH, 200 nM, 16 h) or CDC7i (PHA, 5 µM, 4 h), followed by 250 nM CHK1*i* treatment for 2 h. Cellular TOP1cc levels were detected using anti-TOP1cc monoclonal antibody. (C) RADAR assay in mouse splenocytes treated with CHK1*i*. Freshly isolated mouse splenocytes were treated with 250 nM CHK1*i* with indicated lengths of time, followed by RADAR assay. Scale bars: 10 µm. **** $p<0.0001$.

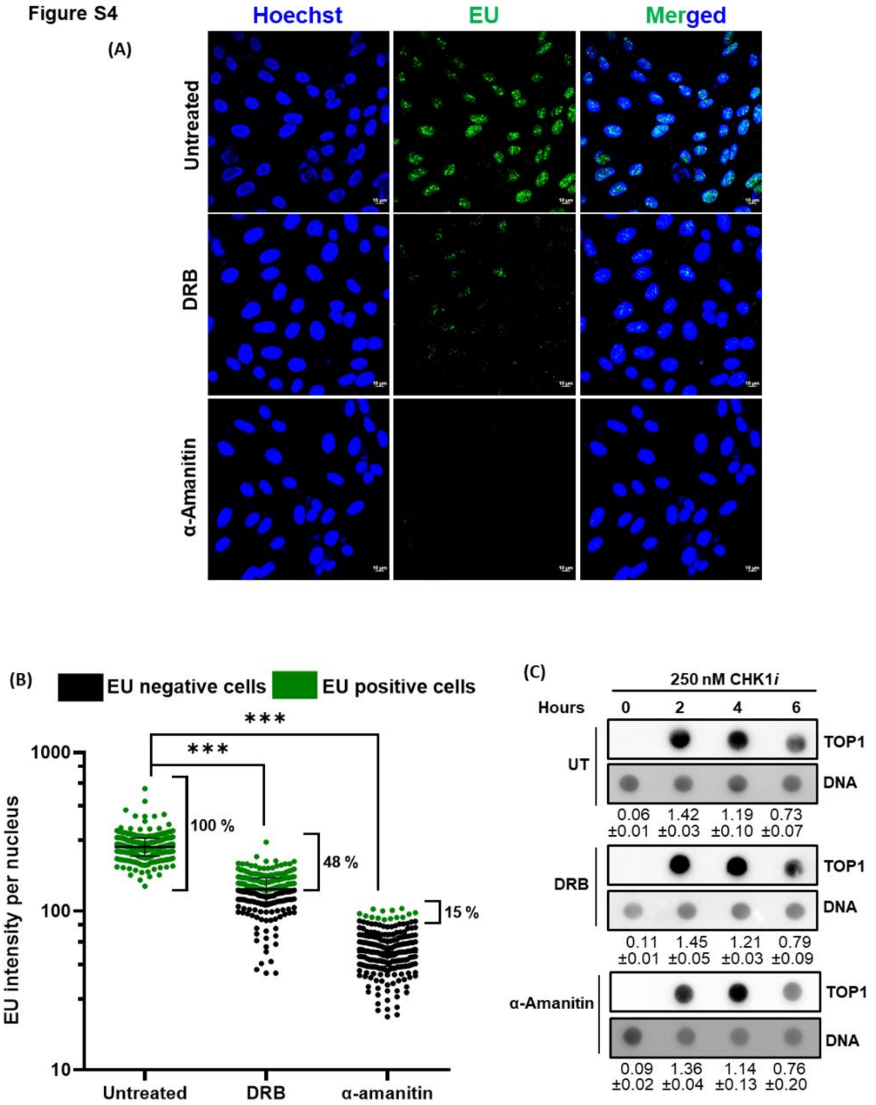


**Figure S4.** Global transcription status in cells treated with the transcription inhibitors DRB and α-Amanitin. U2-OS cells were treated with 100 µM DRB for 4 h or 40 µg/mL α-Amanitin for 16 h, prior to incubation with 200 µM EU for 30 mins prior to termination of experiment.

Global transcription levels were visualized employing click chemistry. Percentage of EU positive cells is displayed along each sample on the graph. (C) TOP1cc stabilization by CHK1*i* in U2-OS cells pre-treated (as mentioned above) with DRB or α-Amanitin. Scale bars: 10 μm. *** $p<0.0001$.

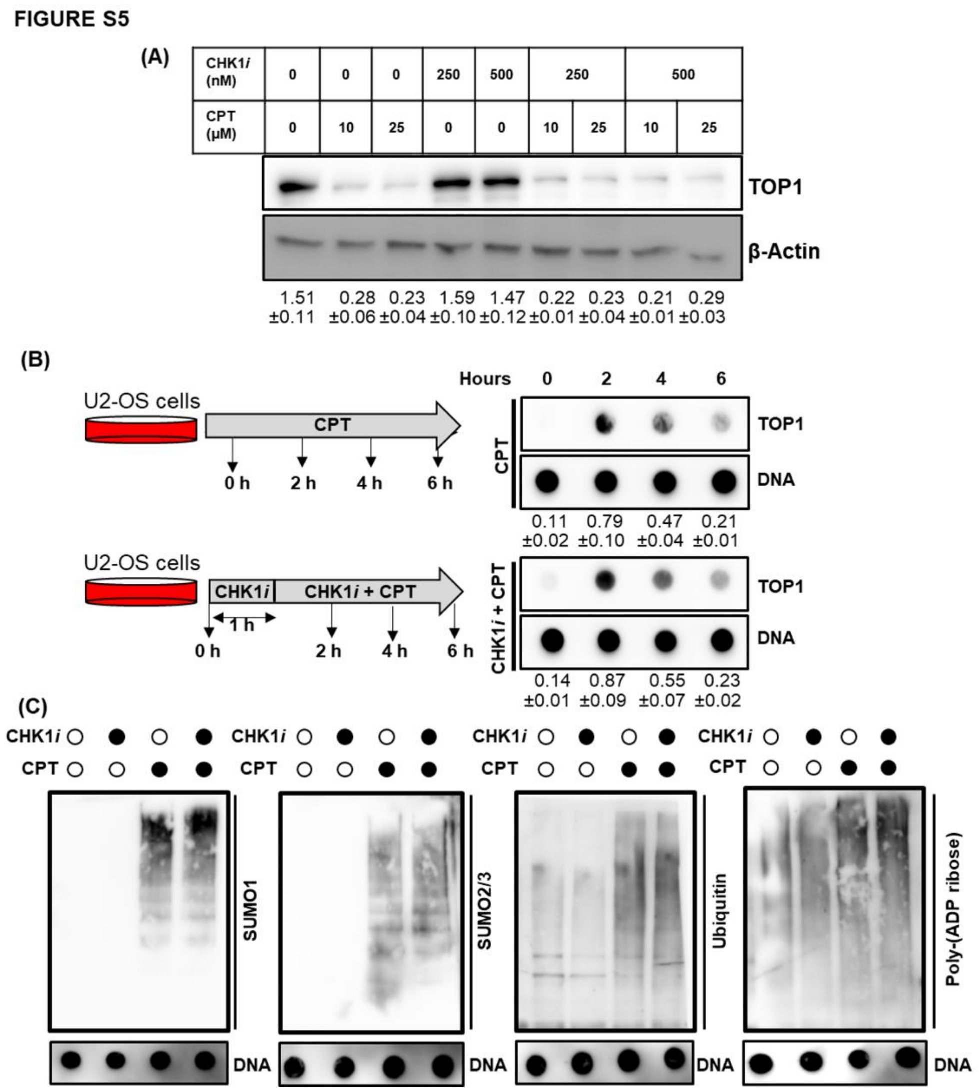


**Figure S5. CHK1 inhibition induced TOP1cc is not readily recognised by clearance machinery.** (A) TOP1 downregulation assay in U2-OS cells treated with CPT with or without pre-treatment with CHK1*i*. U2-OS cells were treated with indicated concentrations of CPT for 1 h with or without 1 h pre-treatment with 250 or 500 nM CHK1*i*. Global TOP1 downregulation was then evaluated by the alkaline lysis method. (B) RADAR assay with U2-OS cells treated with CPT with or without CHK1*i* pre-treatment. U2-OS cells were treated with 1 μM CPT with or without 1 h pre-treatment with 250 nM CHK1*i*, followed by RADAR assay. (C) DUST assay in U2-OS cells treated with 10 µM CPT with or without pre-treatment with CHK1*i*. U2-OS cells were treated with 10 µM CPT for 30 min with or without 1 h pre-treatment with 250 nM CHK1*i*, followed by DUST assay.

**FIGURE S6**

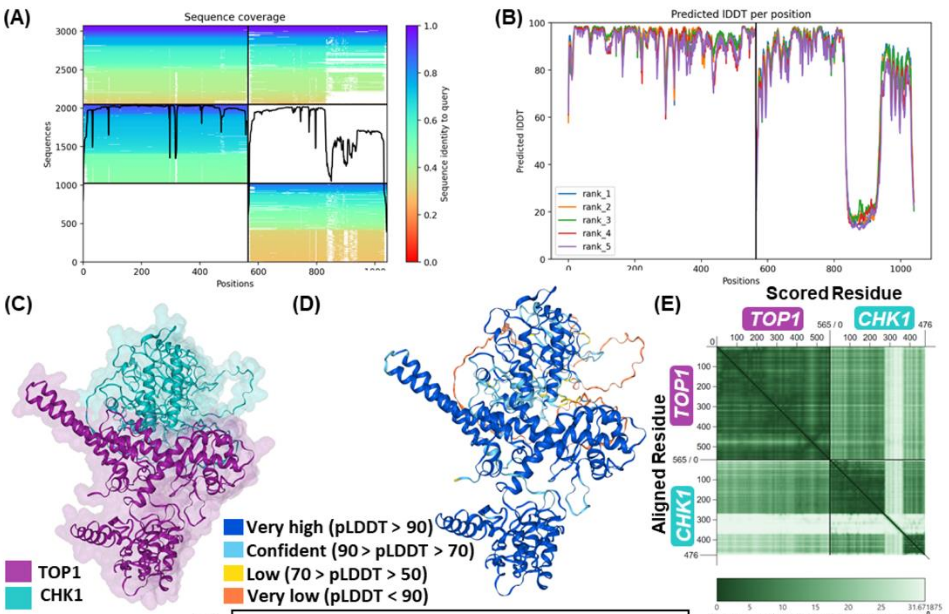


(F)

| Confidence Metrics | | |
|---|---|---|
| Mean pLDDT | pTM | ipTM |
| 84.0622 | 0.7200 | 0.6100 |

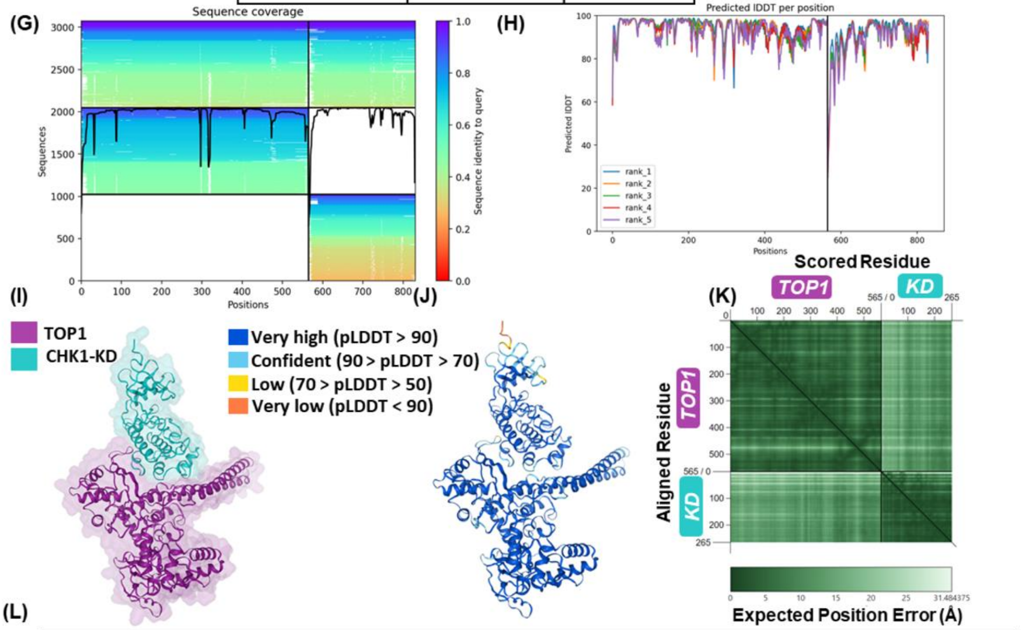


(L)

| Confidence Metrics | | | | | |
|---|---|---|---|---|---|
| Mean pLDDT | pTM | ipTM | SPOC Score | pDOCKQ | Mean PAE |
| 94.2633 | 0.8200 | 0.6400 | 0.414 | 0.349 | 7.5 |

**FIGURE S6 continued**

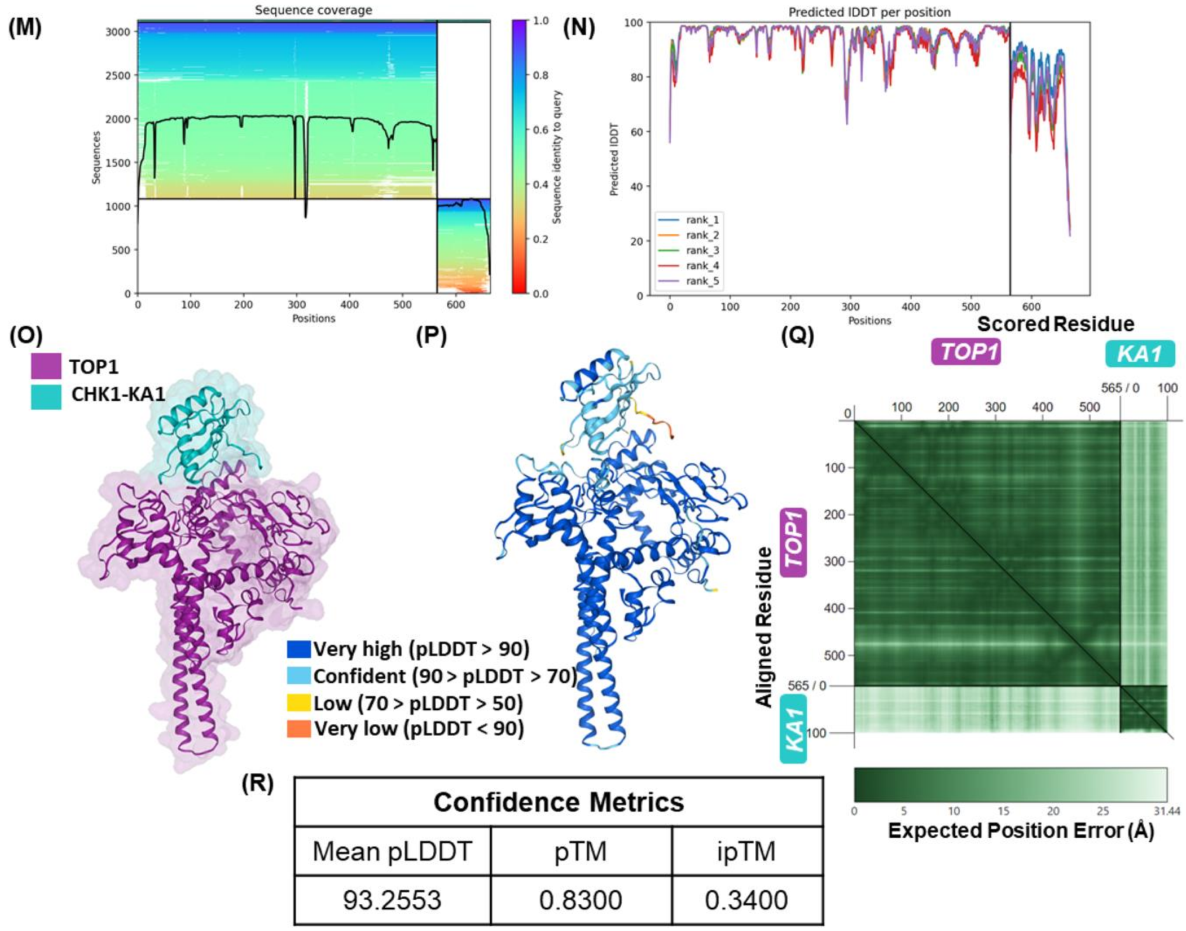


| Confidence Metrics | | |
|---|---|---|
| Mean pLDDT | pTM | ipTM |
| 93.2553 | 0.8300 | 0.3400 |

**Figure S6. Alphafold multimer prediction of TOP1-CHK1 interaction.** Sequence coverage (A) and pLDDT (B) plots of 5 models for TOP1-CHK1 interaction produced by Alphafold Multimer V3. The top ranked model has been reproduced in Figure 2. The second ranked model has been shown in (C) and (D). (E, F) PAE plot and confidence metrics of the second ranked model. Sequence coverage (G) and pLDDT (H) plots of 5 models for interaction between TOP1 and CHK1 kinase domain (KD) produced by Alphafold Multimer V3. (I-J) Visualization of the top-ranked model. (K-L) PAE plot and confidence metrics of the top ranked model. Sequence coverage (M) and pLDDT (N) of 5 models for interaction between TOP1 and CHK1 KA1 domain produced by Alphafold Multimer V3. (O-P) Visualization of the top-ranked model. (Q, R) PAE plot and confidence metrics of the top-ranked model.

FIGURE S7

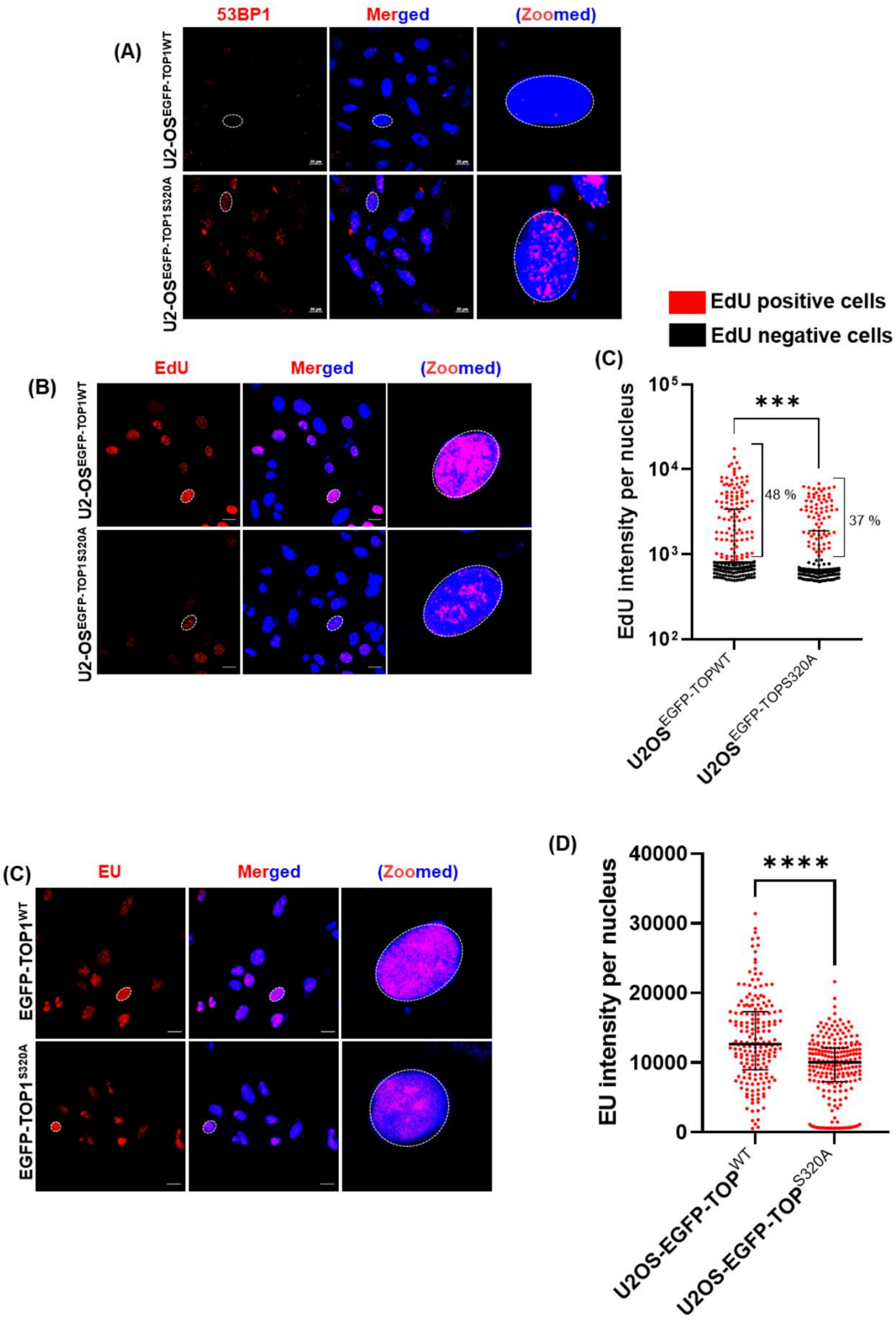


**Figure S7.** (A) Representative images of immunofluorescence imaging of 53BP1 foci in U2-

OS$^{EGFP\text{-}TOP1WT}$ and U2-OS$^{EGFP\text{-}TOP1S320A}$ cells. Quantitative analysis has been shown in Figure 5C. (B) EdU incorporation assay in U2-OS$^{EGFP\text{-}TOP1WT}$ and U2-OS$^{EGFP\text{-}TOP1S320A}$ cells. Cells were labelled with 10 µM EdU for 30 minutes, followed by detection through click chemistry. (C, D) EU incorporation assay in U2-OS$^{EGFP\text{-}TOP1WT}$ and U2-OS$^{EGFP\text{-}TOP1S320A}$ cells. Cells were labelled with 1 mM EU for 1 h, followed by detection through click chemistry. Scale bars 10 µm. **** $p$<0.0001.

**SUPPLEMENTARY TABLE 1**

| INHIBITOR | TARGET KINASE (S) | CONCENTRATIONS |
|---|---|---|
| KU-55933 | ATM | 2.5 µM, 5 µM, 10 µM |
| PV1019 | CHK2 | 2.5 µM, 5 µM, 10 µM |
| AZD6738 | ATR | 500 nM, 1 uM, 2.5 uM |
| SCH900776 | CHK1 | 100 nM, 250 nM, 500 nM |
| NU7026 | DNAPK | 2.5 µM, 5 µM, 10 µM |
| Volasertib (BI627) | PLK1, PLK2, PLK3 | 25 nM, 50 nM, 100 nM |
| Wortmannin | PI3K | 500 nM, 1 µM, 2. 5 µM |
| RO-3306 | CDK1 | 1 µM, 2.5 µM, 5 µM |
| SP600125 | JNK | 5 µM, 10 µM, 20 µM |
| U0126 | MEK1, MEK2 | 2.5 µM, 5 µM, 10 µM |
| Danusertib | Aurora A, Aurora B, Aurora C | 25 nM, 50 nM, 100 nM |
| KN-93 | CAMK | 5 µM, 10 µM, 20 µM |
| LJH685 | RSK1, RSK2, RSK3 | 1 µM, 2.5 µM, 5 µM |
| PHA-767491 | Cdc7 | 2.5 µM, 5 µM, 10 µM |
| GSK2606414 | PERK | 250 nM, 500 nM, 1 µM |
| MRT68921 | ULK1, ULK2 | 250 nM, 500 nM, 1 µM |
| SB203580 | p38 | 2.5 µM, 5 µM, 10 µM |
| ARQ-092 | Akt1, Akt2, Akt3 | 2.5 µM, 5 µM, 10 µM |
| Go6983 | PKCα, PKCβ, PKCγ, PKCδ, PKCζ, PKCµ | 2.5 µM, 5 µM, 10 µM |
| CX-4945 | CK2 | 2.5 µM, 5 µM, 10 µM |
| Brigatinib (AP26113) | ALK | 25 nM, 50nM, 100 nM |
| MK1775 | Wee1 | 50 nM, 100 nM, 200 nM |
| Dorsomorphin (Compound C) | AMPK | 5 µM, 10 µM, 20 µM |
| Torin1 | mTORC1 | 100 nM, 250 nM, 500 nM |
| Roscovitine (Seliciclib) | CDK2, CDK7, CDK9 | 5 µM, 10 µM, 20 µM |

**Supplementary Table 1:** Details of kinase inhibitors, their targets, and concentrations employed in the screening if current investigation.

### SUPPLEMENTARY TABLE 2

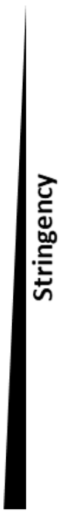


| | Query Motif | Hits | |
|---|---|---|---|
| | | Position | Sequence |
| Minimal Sequence | [RK]-x-x-[ST] | 44 – 47<br>64 - 67<br>70 – 73<br>151 – 154<br>284 – 287<br>310 – 313<br>317 – 320<br>391 – 394<br>443 – 446<br>567 – 570<br>603 – 606<br>615 – 618 | KDREKSKH<br>KEKEKTKH<br>KHKDGSSE<br>PKKIKTED<br>WRKEMTNE<br>YFKAQTEA<br>ARKQMSKE<br>DAKVPSPP<br>WQKYETAR<br>FDRLNTGI<br>QLKELTAP<br>PAKILSYN |
| | [KRHV]-[RK]-x-x-[ST] | 69 – 73<br>150 – 154<br>283 – 287<br>316 – 320 | TKHKDGSSE<br>KPKKIKTED<br>DWRKEMTNE<br>EARKQMSKE |
| Full Sequence | [RFPGAILV]-[KRHV]-[RK]-x-x-[ST] | 149 – 154<br>315 – 320 | YKPKKIKTED<br>TEARKQMSKE |

**Supplementary Table 2:** Prediction of CHK1 target motifs on TOP1. While the entire motif is highlighted in green, the target residue is depicted in red.

SUPPLEMENTARY TABLE 3

| Site | Untreated | 250 nM SCH 2h | 250 nM SCH 6h | 200 nM CPT 2h | 1 μM CPT 2h | Previously Detected? |
|---|---|---|---|---|---|---|
| S320 | + | + | + | + | + | No |
| S394 | + | + | - | - | + | Yes[23,61] |
| T570 | + | - | - | + | - | Yes[61] |
| Y231 | + | - | - | - | - | No |
| S250 | + | - | - | + | - | No |
| Y480 | + | - | - | + | - | Yes[61] |
| T446 | - | + | - | - | - | No |
| Y461 | - | - | + | + | - | No |
| S534 | - | - | - | + | - | No |
| Y538 | - | - | - | + | - | No |
| T706 | - | - | - | + | - | Yes[61] |

**Supplementary Table 3:** Details of sites found to be phosphorylated on catalytically active TOP1. Whether they have previously been reported elsewhere is also indicated with references[23,61].

**SUPPLEMENTARY TABLE 4**

| | Forward Primer | Reverse Primer |
|---|---|---|
| **MYO3A TSS** | GTCAGATCCGGAGGACC | GTTCATCCCTCTCCTCCC |
| **MYO3A Gene Body** | CTACAGCAGCCCACTCAAG | CATTATCCAGTTTCTTGATTCAT G |
| **β-Actin TSS** | CGGGGTCTTTGTCTGAGC | CAGTTAGCGCCCAAAGGAC |
| **β-Actin Gene Body** | GGAGCTGTCACATCCAGGGTC | TGCTGATCCACATCTGCTGG |
| **g-Actin TSS** | CCGCAGTGCAGACTTCCGAG | CGGGCGCGTCTGTAACACGG |
| **g-Actin Gene Body** | GTGACACAGCATCACTAAGG | ACAGCACCGTGTTGGCGT |
| **PTB Gene Body** | GCCGTTGGTACAAAGGTAGG | GCCCCTTAGGAATGGAAAAG |
| **Geminin 7 Gene Body** | TCTTCTTCCACCTGGACCAC | GGGACAGAGAGAGTGCCTTG |
| **IL4 Gene Body** | TTCAGGTGACAAGTGCCACAG | CTGGTTGGCTTCCTTCACAG |
| **Intergenic** | ACCCAGCACCCCCTAATACC | AGCCGGACATGCTTCCAGAG |

**Supplementary Table 4:** Details of primers used in qPCR.
